# Estrogen Receptor Enhancers Sensitive to Low Doses of Hormone Specify Distinct Molecular and Biological Outcomes

**DOI:** 10.1101/2025.08.08.669412

**Authors:** Hyung Bum Kim, Tulip Nandu, Cristel V. Camacho, W. Lee Kraus

## Abstract

Adult women are typically exposed to estradiol (E2) concentrations of ∼100-200 pM, yet most cell-based studies use 100 nM. We determined the molecular effects of E2 concentrations spanning six orders of magnitude (1 pM to 100 nM) in breast cancer cells. Estrogen receptor alpha (ERα) enhancers formed at low physiological doses of E2 (1-100 pM) are mechanistically distinct from those that form at high pharmacological doses (10-100 nM). They (1) form in open chromatin bound by FOXA1, (2) produce enhancer RNAs enriched with functional eRNA regulatory motifs (FERMs), and (3) drive expression of cell proliferation genes with promoter-proximal paused RNA polymerase II. Importantly, low dose ERα enhancer usage is elevated in breast cancer patients with poor responses to aromatase inhibitors, likely as a continued response to low circulating levels of E2. Collectively, our results identify mechanistic differences between low and high dose ERα enhancers that specify distinct biological outcomes.

## Introduction

Estrogens are a class of steroid hormones required for the normal development and function of reproductive organs, mammary glands, bone, heart, vasculature, adipose, and key cell types in the central nervous system (Chen et al., 2022; Hamilton et al., 2017; Hewitt et al., 2005; Kronenberg et al., 2007; Pakdel, 2023). Estrogens also play key roles in diseases associated with the same tissues and contribute to a host of pathologies with unique onset, frequency, and presentation in women, such as cardiovascular, neuronal, and immune disease, as well as aging and cancer (Burns and Korach, 2012; Chen et al., 2022; Hamilton et al., 2017; Isola et al., 2023; Pakdel, 2023). Within an individual, estrogen levels may fluctuate dramatically and acutely across multiple time scales: hours, days, months, years, and decades, and can be influenced by physiology, pathology, and therapy [(Jett et al., 2022; Matyi et al., 2019; Pesce et al., 2023) and references therein]. The circulating blood levels of 17β-estradiol (E2), the most potent naturally-occurring estrogen, fluctuate depending on life stage, from non-pregnant lows of ∼70 pM to highs of ∼1.8 nM, rarely exceeding these levels (prepubertal ∼70 pM; menstrual ∼220 pM to 1.8 nM; pregnant 8 nM to 50 nM; and postmenopausal ∼0 to 110 pM) (Carmina et al., 2013; Holst et al., 2004) (Fig. 1A). Despite this understanding, there is gap in our knowledge about the effect of dose on the cellular, molecular, and genomic effects of estrogens.

**Figure 1.**
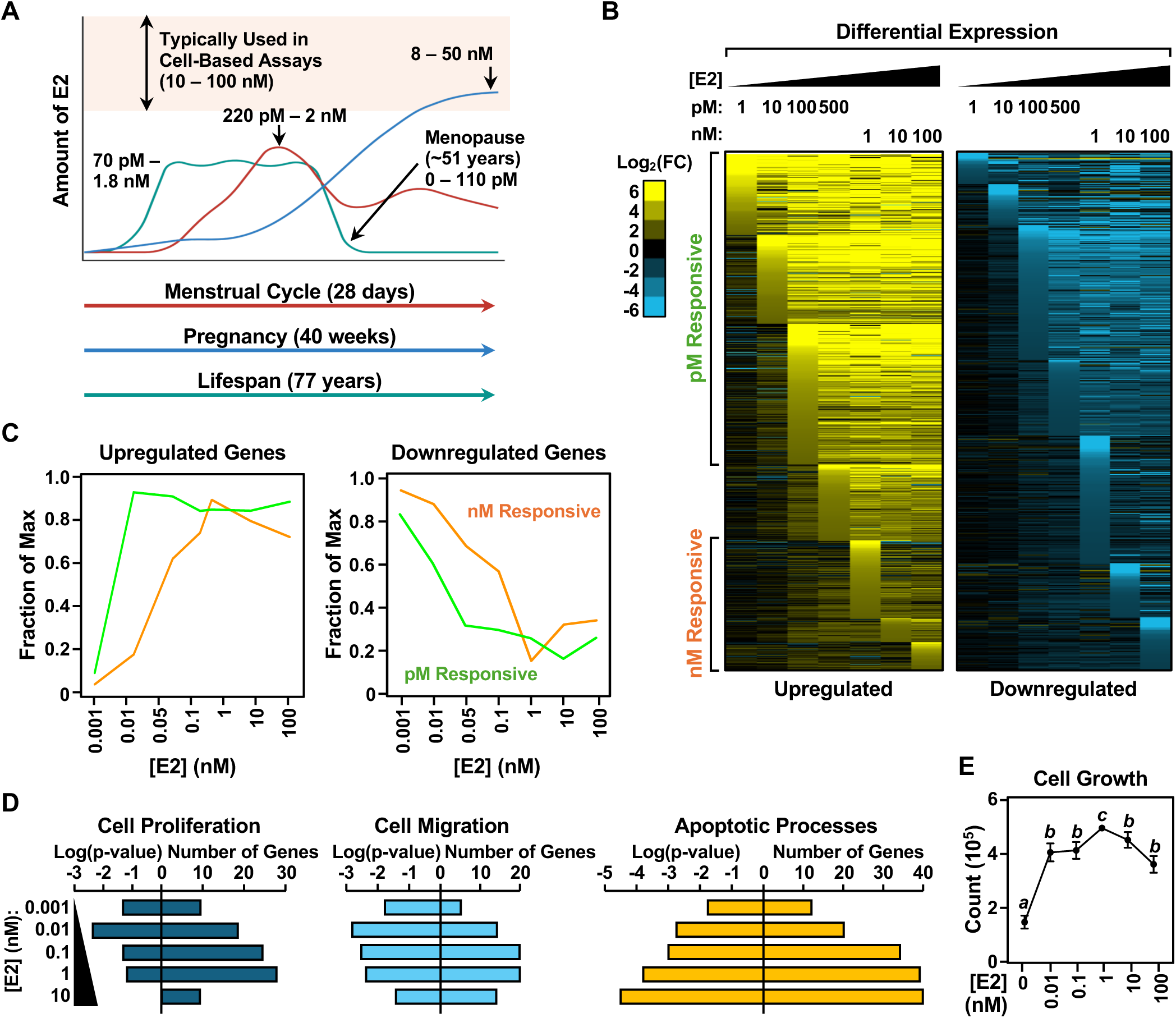
Dose-dependent effects of E2 on ERα-mediated gene expression. **(A)** Schematic diagram showing the ranges of physiological E2 concentrations in women during the mensural cycle, pregnancy, and lifetime. Highlighted region shows the concentrations typically used in biochemical assays. **(B)** Heatmap representations of gene expression as determined by RNA-seq after treatment with increasing concentrations of E2 (1 pM to 100 nM) for 2 hours showing upregulated genes (*left*) and downregulated genes (*right*). Genes are clustered by the dose at which the first significant response occurred (p-value >0.01) in descending order of fold change within the cluster. Gene groups are labeled in aggregate as pM responsive (1 pM to 100 pM E2) and nM responsive (1 nM to 100 nM E2). **(C)** Line graphs showing the relative expression of genes (fraction of maximum) within each dosage group for upregulated (*left*) and downregulated (*right*) over a range of E2 concentrations. **(D)** Gene ontology (GO) analyses showing the number of genes and p-values for the terms cell proliferation, cell migration, and apoptotic process over a range of E2 concentrations. **(E)** Line graph showing cell growth at day 5 of E2 treatment as determined by live cell counts over a range of E2 concentrations. Each point represents the mean ± SEM; n = 3; Points marked with different letters are significantly different from each other. Student’s t-test, p < 0.005.

Many of the biological actions of estrogens are mediated by estrogen receptor alpha (ERα), a nuclear receptor, which functions primarily as a ligand-regulated, DNA-binding transcription factor (TF) in the nuclei of estrogen-responsive cells (Couse and Korach, 1999; Warner et al., 1999; Welboren et al., 2009). Studies of nuclear receptors have played a key role in our understanding of enhancers, including key concepts such as nuclease (e.g., DNase I) hypersensitivity (Hecht et al., 1988; Jantzen et al., 1987; Nyborg and Spindler, 1986; Usala et al., 1988; Zaret and Yamamoto, 1984), coregulators (Acevedo and Kraus, 2004; Biddie et al., 2010; Glass and Rosenfeld, 2000; Lonard and O’Malley, 2012; McKenna et al., 1999), distal enhancers, and chromatin looping (Cheung and Kraus, 2010; Martens et al., 2011). Estrogens act as potent mitogens in ERα-positive breast cancers (ER+ BC), eliciting a rapid and robust genomic response through ERα. Expression of nearly half of the genome is regulated by E2 stimulation in ER+ MCF-7 breast cancer cells (Hah et al., 2011; Hah et al., 2013), making them a powerful system to study the genomic actions of hormonal signaling.

ERα is a transcription factor that resides in the nuclei of cells even in the absence of estrogen. It dimerizes after binding its ligands, including the predominant naturally-occurring estrogen 17β-estradiol (E2), and binds to many thousands of ERα binding sites (ERBSs) across the genome, collectively called the ERα ‘<u>cistrome</u>’ (Carroll et al., 2005; Carroll et al., 2006; Hewitt et al., 2012; Lin et al., 2007; Welboren et al., 2009). ERα binding sites may be pre-bound by ‘pioneer’ transcription factors (e.g., FOXA1) prior to estrogen treatment, which facilitate the binding of ERα to chromatin (Hurtado et al., 2011; Tan et al., 2011). Many ERα binding sites contain a DNA sequence motif called the estrogen response element (ERE) (Warner et al., 1999). The binding of ERα to genomic DNA promotes the coordinated recruitment of coregulator proteins (e.g., NCOAs) that establish active enhancers, leading to chromatin looping and target gene transcription (Foulds et al., 2013; Fullwood et al., 2009; Hah et al., 2013; He et al., 2012; Shlyueva et al., 2014). Enrichment of molecular features, including certain histone modifications (e.g., H3 lysine 27 acetyl, H3K27ac), coactivators (e.g., p300/CBP, Mediator), RNA polymerase II (Pol II), an open chromatin architecture (e.g., DNase I hypersensitivity), and looping to target gene promoters, mark or identify active enhancers (Bulger and Groudine, 2011; Hah et al., 2013; Melgar et al., 2011; Natoli and Andrau, 2012). The transcriptional effects of estrogen are rapid, on the order of minutes, resulting in transcription at both target genes and their enhancers (Hah et al., 2011; Hah and Kraus, 2014; Hah et al., 2013; Hou and Kraus, 2021).

Although the importance of dose, and the frequency and duration of administration, have long been recognized as important aspects of the therapeutic use of steroid hormones (Ansbacher, 2000; Cho et al., 2023; Paszkowski et al., 2019), the effects of lower physiological doses of E2 and acute treatment on the molecular and genomic endpoints listed above are not well characterized. In fact, previous genomic studies of nuclear receptor function have relied nearly exclusively on pharmacologic doses of hormone. Here, we describe studies examining the dose effects of E2 for concentrations spanning six orders of magnitude (1 pM to 100 nM) in MCF-7 cells. We found that ERα enhancers formed at low (pM) physiological doses are mechanistically and functionally distinct from those that form at high (nM) pharmacological doses.

## Results

### Different E2 dose sensitivities for the expression of distinct sets of estrogen target genes

To determine how ERα-expressing cells respond to different doses of E2, we treated MCF-7 human breast cancer cells with a range of physiological to pharmacological doses of E2 over six orders of magnitude (1 pM to 100 nM) and examined the changes in gene expression by RNA-seq. Remarkably, we observed upregulation or downregulation of distinct sets of genes even at the lowest doses of E2 (Fig. 1B). At each successive dose, additional regulated genes were added to the previous genes sets (Fig. 1B). This analysis defined distinct sets of genes responsive to lower, physiological concentrations of E2 (“pM responsive genes”) or higher, pharmacological concentrations of E2 (“nM responsive genes”) (Fig. S1, A and B), with a clear dose shift evident for the different gene sets when expressed in aggregate (Fig. 1C).

Pathway analysis using the Kyoto Encyclopedia of Genes and Genomes (KEGG) showed enrichment of a number of terms related to cancer pathology in the pM responsive gene group, including DNA replication, Wnt signaling, basal-cell carcinoma, and proteoglycans in cancer (Fig. S1C). In contrast, a thematically broader range of terms, including cellular senescence, insulin signaling, MAPK signaling, and tight junctions, was enriched in the nM responsive group (Fig. S1D). Gene ontology (GO) analyses showed dose-dependent enrichment of cancer-related terms, with cell proliferation and cell migration showing greater enrichment with increasing concentrations of E2, except at the highest dose (100 nM) (Fig. 1D). Cell growth assays across a range of E2 concentrations showed a corresponding trend, with the highest dose (100 nM) eliciting a submaximal response compared to 1 nM E2 (Fig. 1E). Taken together, these analyses reveal key features of the E2 dose response: (1) E2 is a potent regulator of ERα-dependent gene expression and cell growth, with robust responses occurring at pM concentrations of E2 and (2) E2 responses saturate at nM concentrations of E2, with the highest doses not necessarily eliciting the greatest responses.

### Different chromatin states at pM responsive and nM responsive ER**α** enhancers

To better understand the molecular mechanisms underlying the dose-dependent formation of ERα enhancers that direct the expression of estrogen target genes in response to E2, we determined the ERα cistrome in MCF-7 cells across a range of E2 concentrations by ChIP-seq (Fig. 2A; Fig. S2A). We observed that a subset of the ERα cistrome formed at low (pM) concentrations of E2 and expanded at higher doses, with the sites of ERα binding at lower doses being a subset of those observed at higher doses (i.e., the ERα binding sites at higher doses encompasses nearly all of those observed at lower doses; Fig. S2, A and B). Interestingly, ERα bound the ‘low dose’ enhancers at 10 pM E2 to the same extent that it bound to the ‘high dose’ enhancers at 10 nM E2 based on average read counts for each class (Fig. 2, B and C). Based on these analyses, we also defined the doses at which genomic sites first became accessible to ERα binding.

**Figure 2.**
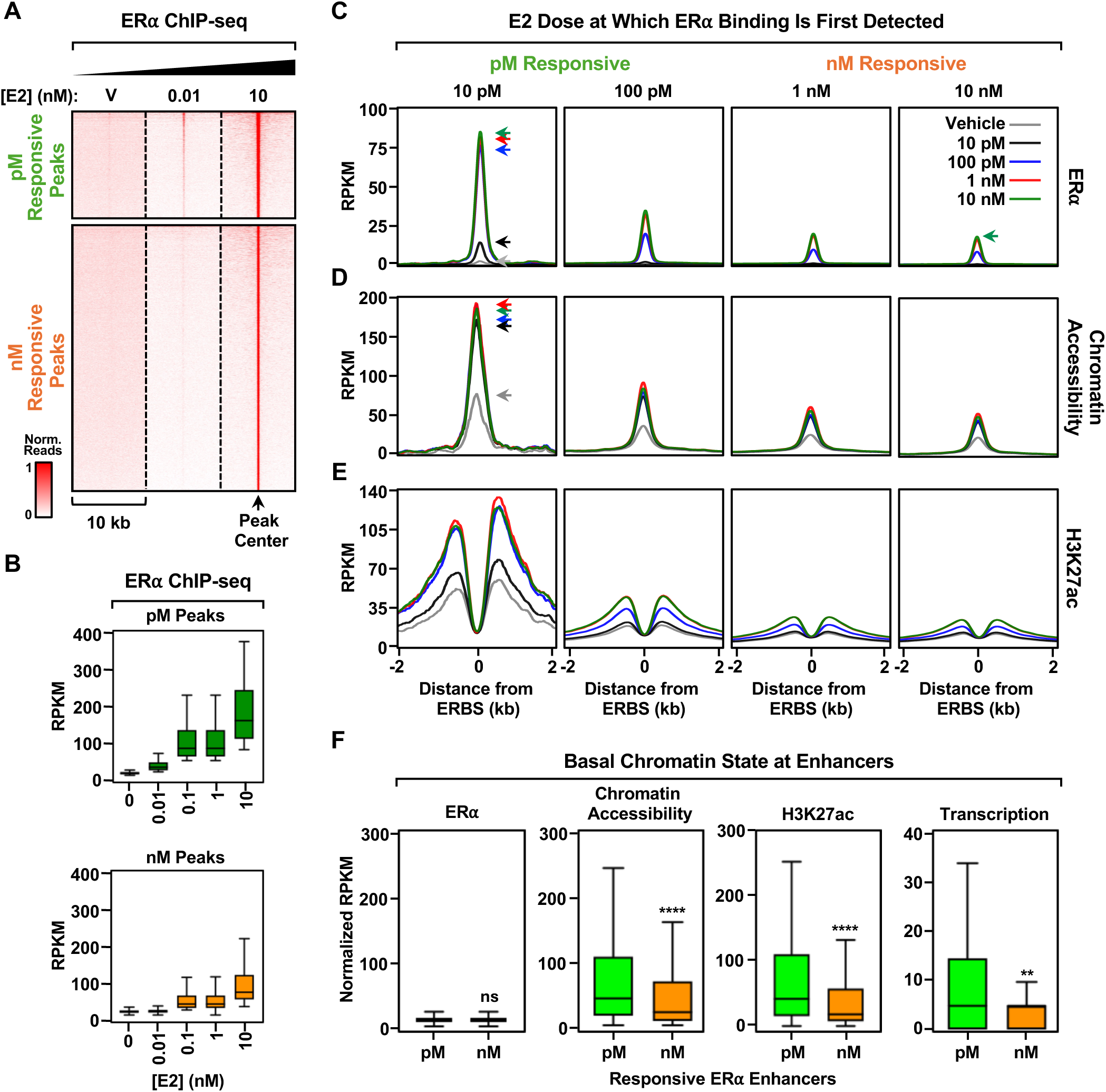
Analysis of chromatin state at pM responsive and nM responsive ERα enhancers. **(A)** Heatmap showing ERα binding across the genome as determined by ChIP-seq after 60 minutes of treatment with vehicle (DMSO), 10 pM E2, or 10 nM E2. ERBS groups are labeled in aggregate as pM peaks (1 pM to 100 pM E2) and nM peaks (1 nM to 100 nM E2). **(B)** Boxplots showing ERα read counts at pM responsive (*top*) and nM responsive (*bottom*) ERBS for a range of E2 concentrations. **(C through E)** Metaplots showing the read counts in RPKM around ERBS from genomic assays over a range of E2 concentrations (10 pM to 10 nM). (C)_ERα binding by ChIP-seq, (D) chromatin accessibility by ATAC-seq, and (E) H3K27ac enrichment by ChIP-seq. Peaks are centered on the summits of significant ERBSs. **(F)** Boxplots showing the normalized read counts for ERα binding (ChIP-seq), chromatin accessibility (ATAC-seq), H3K27ac enrichment (ChIP-seq), and transcriptional activity (PRO-seq) at pM responsive and nM responsive ERBSs prior to E2 treatment. Wilcoxon Rank Sum test, ns = not significant, ** p < 5 × 10^-10^, **** p < 5 × 10^-15^.

In subsequent analyses, we assayed additional features associated with ERα binding sites, including chromatin accessibility (by ATAC-seq) and H3K27ac enrichment (by ChIP-seq) (Fig. 2, D and E; Fig. S2, C and D). As with ERα binding, chromatin accessibility and H3K27ac enrichment were greatest at sites that first exhibited ERα binding at the lowest doses of E2 (Fig. 2, D and E). Moreover, we observed that the low dose (pM) ERα binding sites were more open, accessible, and active in the basal state (without E2 treatment) than the high dose (nM) sites (Fig. 2F; note: for these analyses, combined data from the 10 and 100 pM E2 treatments groups are designated pM* and combined data from the 1 and 10 nM E2 treatments groups are designated nM*). These analyses included enhancer transcription data from PRO-seq (Fig. 2F, *right*); enhancer transcription is a robust indicator of enhancer activity (Hah et al., 2013; Hou and Kraus, 2021). Taken together, these results demonstrate that enhancers that form at ERα binding sites in response to low doses of E2 do so at accessible sites in the genome.

### Different FOXA1 and GATA3 usage at pM responsive and nM responsive ER**α** enhancers

The efficient formation of ERα enhancers requires cooperation with FOXA1 and GATA3, which have been described as ‘pioneer factors’ for ERα (Carroll et al., 2005; Carroll et al., 2006; Hurtado et al., 2011; Martin et al., 2021; Takaku et al., 2020; Theodorou et al., 2013). To determine if pM responsive and nM responsive ERα enhancers might use FOXA1 and GATA3 differently, we performed a de novo motif analysis in the vicinity of ERα binding sites. We observed a significant enrichment of the canonical palindromic estrogen response element (ERE), with the greatest enrichment in the enhancers established at 10 pM E2 (Fig. 3, A through C). A similar analysis revealed significant enrichment of FOXA1 motifs at ERα binding sites, also with the greatest enrichment in the enhancers established at 10 pM E2, but with a precipitous reduction for enhancers formed at higher doses (Fig. 3, A through C). In contrast, GATA3 motifs showed the greatest enrichment at ERα binding sites established at higher doses of E2 (Fig. 3, A through C).

**Figure 3.**
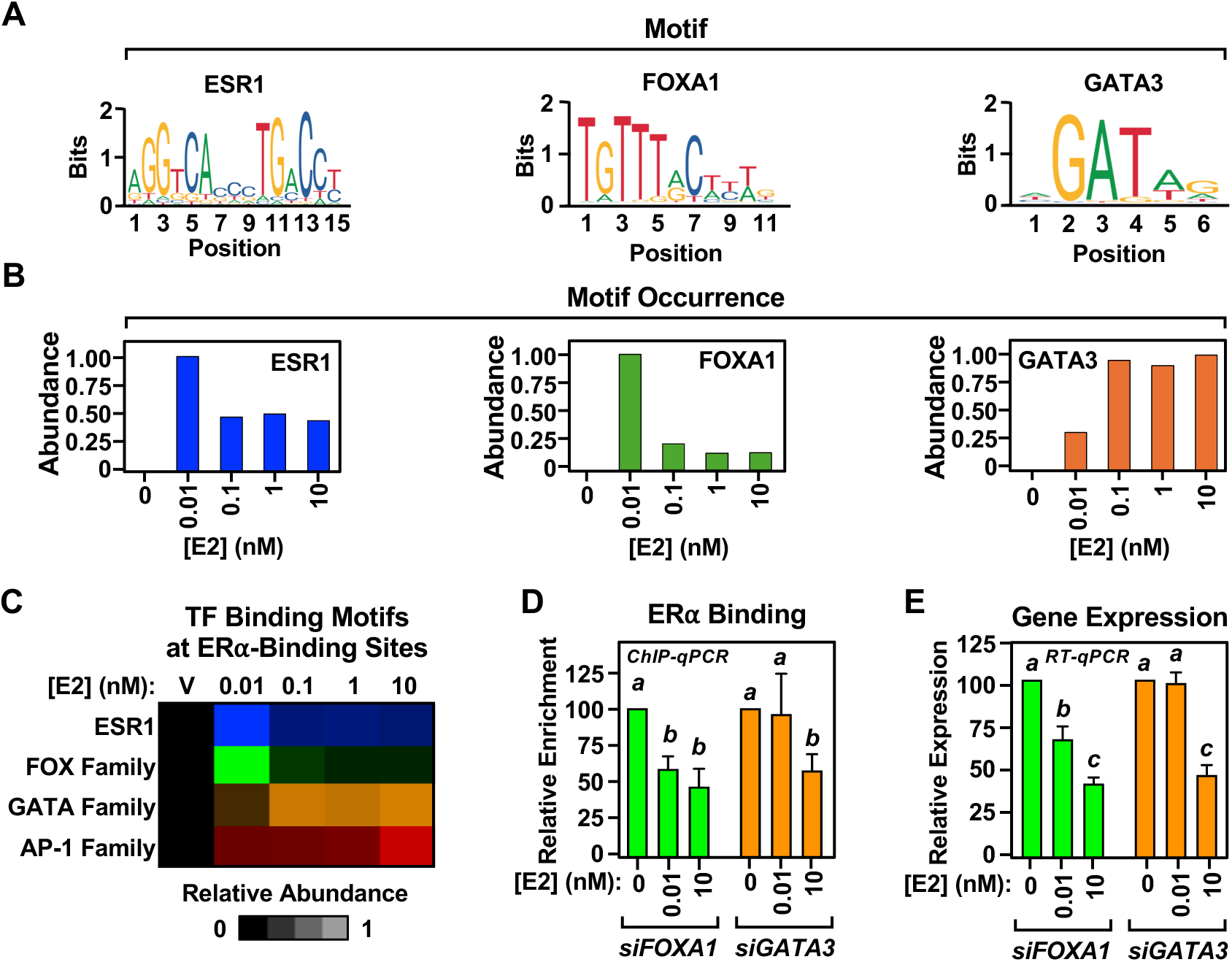
Dose-dependent enrichment of FOXA1 and GATA3 motifs near ERα binding sites. **(A)** JASPAR gene logos of ESR1, FOXA1, and GATA3 motifs found ± 200 bp around ERBSs. **(B)** Bar graphs showing the abundance of motif occurrences around ERBSs established over a range of E2 concentrations, normalized to their respective maximum values. **(C)** Heatmap showing the relative occurrences of ESR1, FOX family (FOXA1, FOXA2, FOXO1), GATA family (GATA3, GATA4, GATA6), and AP-1 family (FOS, c-JUN, ATF2, ATF3) motifs over a range of E2 concentrations, normalized to their respective maximum values. **(D and E)** Effects of FOXA1 or GATA3 depletion by siRNA-mediated knockdown on (D) ERα binding to chromatin after 60 minutes of E2 treatment and (E) E2-dependent gene expression after 2 hours of E2 treatment. Bar graphs showing (D) relative enrichment from ERα ChIP-qPCR and (E) gene expression from RT-qPCR for E2-responsive enhancers and their cognate target genes. The data are expressed in aggregate for a collection of ERBS and target genes (*TFAP2C*, *TSKU*, *GREB1*, *MBOAT1*, *MIDEAS*, *CYP1B1*, *WNT16*, *ADRB1*, *PGR*). The data are normalized to the control siRNA. Each bar represents the mean + SEM; n = 3. Bars marked with different letters are significantly different from each other. Student’s t-test, p < 0.005.

To explore this in more detail, we examined the binding of ERα to bulk chromatin at different doses of E2 with or without siRNA-mediated knockdown of *FOXA1* and *GATA3*. Depletion of FOXA1 and GATA3 was confirmed by Western blotting (Fig. S3A). As with the motif enrichment, we observed a greater enrichment of FOXA1 in bulk chromatin at 10 pM E2 than at 10 nM E2, and vice versa for GATA3 (Fig. S3A). As expected, depletion of FOXA1 or GATA3 impaired ERα binding, with the effects of GATA3 most pronounced at 10 nM E2 (Fig. S3A). Similar results were observed in locus-specific ChIP-qPCR assays for a collection of nine ERα enhancers and their target genes, expressed in aggregate (Fig. 3, D and E).

Additional genome-wide assays reinforced this conclusion. FOXA1 and GATA3 ChIP-seq at 10 pM and 10 nM E2 showed thousands of significant peaks for both factors across the genome in both conditions, many of which overlapped ERα binding sites (Fig. 4A; Fig. S3B). Treatment with 10 pM E2 promoted the greatest enrichment of FOXA1 at ERα binding sites, whereas 10 nM E2 promoted the greatest enrichment of GATA3 (Fig. 4B; Fig. S3C), with a significantly greater GATA3/FOXA1 ratio for the nM* dosage group (Fig. S3D). The expansion of GATA3 across the ERα cistrome at the highest doses of E2 was evident in a Venn diagram plotted versus ERα binding at the various doses (Fig. 4C). Finally, we used rapid immunoprecipitation mass spectrometry of endogenous proteins [RIME (Mohammed et al., 2016); a.k.a., ChIP mass spectrometry] as an orthogonal approach to monitor the dose-dependent enrichment of FOXA1 and GATA3 to chromatin-bound ERα. We observed E2-dependent recruitment of ERα to chromatin, with similar enrichment at both 10 pM and 10 nM E2 (Fig. 4D, *top*). As with the ChIP-seq results, we observed the greatest enrichment of FOXA1 at the lower dose of E2 and the greatest enrichment of GATA3 at the higher dose of E2 (Fig. 4D, *bottom*). Thus, in multiple different assays, we observed similar results.

**Figure 4.**
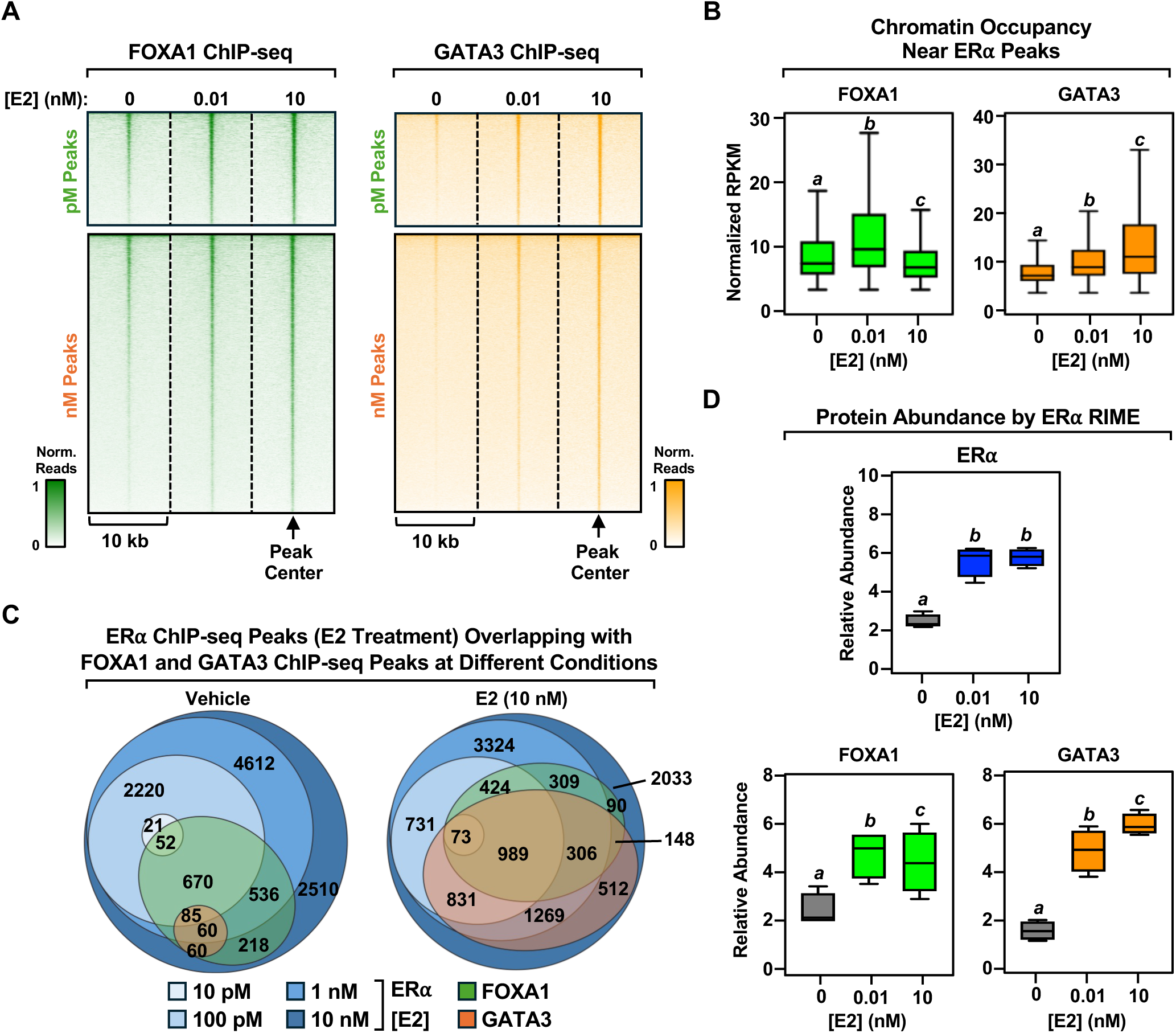
Dose-dependent binding of FOXA1 and GATA3 at ERα binding sites. **(A)** Heatmap showing FOXA1 (*left*) and GATA3 (*right*) chromatin occupancy as determined by ChIP-seq for vehicle (V), 10 pM E2, and 10 nM E2 treatments at pM responsive and nM responsive ERBS. **(B)** Boxplots showing the binding of FOXA1 (*left*) or GATA3 (*right*) at ERBS in response to low (10 pM) and high (10 nM) E2 treatment. Bars marked with different letters are significantly different from each other. Wilcoxon Rank Sum test, p < 5 × 10^-10^. **(C)** Euler diagrams showing overlaps of ERα ChIP-seq peaks across different E2 treatment concentrations (10 pM to 10 nM) with FOXA1 and GATA3 peaks after treatment with vehicle (*left*) or 10 nM E2 (*right*). **(D)** Boxplots showing the relative abundance of FOXA1 and GATA3 associating with chromatin-bound ERα in response to low (10 pM) and high (10 nM) E2 treatment as determined by RIME. The data are normalized to the respective IgG controls. Bars marked with different letters are significantly different from each other. Student’s t-test, p < 0.005.

Collectively, our results indicate different FOXA1 and GATA3 usage at pM responsive and nM responsive ERα enhancers, with FOXA1 required for enhancer formation and target gene expression at low dose enhancers and genes, and GATA3 required at high dose enhancers and genes.

### Different modes of chromatin accessibility mediated by FOXA1 and GATA3 proximal to ER**α** binding sites

To better understand the differential usage of FOXA1 and GATA3 at pM responsive and nM responsive ERα enhancers, we considered the location of EREs relative to open regions of chromatin, as determined by ATAC-seq. We began by inspecting examples of pM responsive and nM responsive enhancers: *XBP1* and *PGR*, which exhibit ERα binding at 10 pM and 10 nM E2, respectively (Fig. 5A). We observed that *XBP1* had four EREs in open regions of chromatin established by FOXA1 prior to E2 treatment, while *PGR* had none (Fig. 5B). The *PGR* enhancer, however, gained two newly available EREs only after treatment with E2. This suggests that ERα binding occurs in the presence of E2 when EREs are available in an accessible region of chromatin; this occurs before E2 treatment for FOXA1 and after E2 treatment for GATA3. This is observed globally, as treatment with E2 increases the total number of EREs in accessible regions across the genome (Fig. 5C).

**Figure 5.**
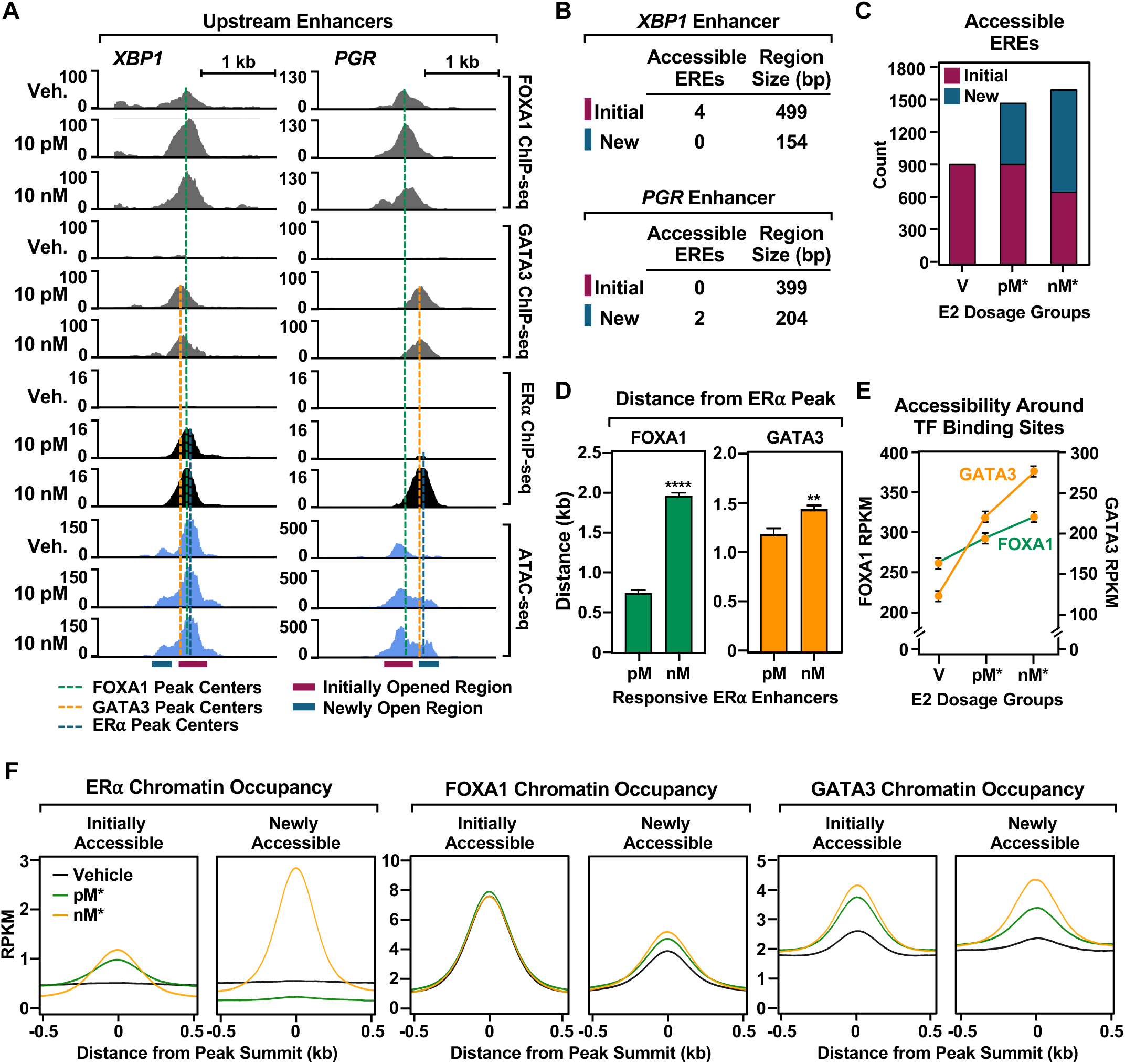
ERα binding at pM responsive enhancers occurs in open chromatin while binding at nM responsive enhancers occurs at newly opened regions. **(A)** Browser tracks for upstream enhancer regions of representative pM responsive (*XBP1*) and nM responsive (*PGR*) genes showing FOXA1, GATA3, and ERα ChIP-seq data with ATAC-seq data. Dotted lines highlight peak summits for FOXA1 (*green*), GATA3 (*orange*), and ERα (*blue*). Highlights below the tracks show regions that are open prior to (*maroon*) or after (*dark blue*) E2 induction as determined by ATAC-seq. **(B)** Table showing the number of ERE occurrences and length (in bp) of initially or newly open regions near the *XBP1* and *PGR* genes from panel (A) as determined by FIMO with a p-value cutoff of < 0.005. **(C)** Bar graphs showing the total number of ERE occurrences in open regions of chromatin that were available prior to (*maroon*) or after (*dark blue*) E2 treatment shown for two different dosage groups (pM = 1, 10, 100 pM; nM = 1, 10, 100 nM). **(D)** Bar graphs showing that average distances between ERα and FOXA1 (*green*) or GATA3 (*orange*) ChIP-seq peak summits within a 2 kb range for pM responsive and nM responsive ERBS. Each bar represents the mean + SEM. Student’s t-test, ** p < 0.001, **** p < 0.0001. **(E)** Line graphs showing chromatin accessibility around FOXA1 or GATA3 peaks from ChIP-seq for two different E2 dosage groups. Each point represents the mean ± SEM. **(F)** Metaplots of ERα, FOXA1, and GATA3 chromatin occupancy as determined by ChIP-seq within regions of accessible chromatin prior to (*left*) and after (*right*) E2 treatment as determined by ATAC-seq.

We also observed that the distance between ERα and FOXA1 binding sites is associated with dose dependency. While ERα binding sites at pM responsive enhancers are generally in close proximity to FOXA1 binding sites (mean of 740 bp), ERα binding sites at nM responsive enhancers are nearly threefold further away from FOXA1 binding sites (mean of 1,915 bps) (Fig. 5D). In contrast, we observed little difference in the distance between ERα binding sites and GATA3 binding sites between pM and nM responsive enhancers (Fig. 5D). Additionally, regions of open chromatin near GATA3 binding sites showed a significantly larger increase in accessibility upon E2 treatment compared to FOXA1 binding sites, which showed a more modest increase in accessibility (Fig. 5E). Next, we investigated patterns of TF binding at initially accessible (prior to E2 treatment) and newly accessible (after E2 treatment) regions. In agreement with the results described above, we found that while ERα bound at initially open regions of chromatin in response to both pM and nM E2 treatments, only nM E2 treatment promoted ERα binding at newly open regions of chromatin (Fig. 5F). This likely occurs because these regions are not yet sufficiently open at pM doses of E2.

Taken together, our results indicate that FOXA1 initially establishes open regions of chromatin accessible to ERα at low (pM) doses of E2, while GATA3 drives newly opened regions of chromatin after E2 induction in proximity to FOXA1 binding sites. These open regions of chromatin facilitate ERα binding if an ERE is present. With E2 treatment, larger regions become accessible, allowing ERα binding at sites farther away from the initial regions established by FOXA1.

### AP-1 actions at ER**α** enhancers occur at high doses of E2

Previous studies have suggested that the transcription factor AP-1 (Fos/Jun heterodimers) may also be enriched at ERα binding sites and may, in some cases, tether ERα to chromatin without direct binding of the receptor to DNA (Carroll et al., 2006; Gao et al., 2008; Kushner et al., 2000; Lin et al., 2007). These analyses were performed with high (>10 nM) E2 treatment. When mining our data, we observed the greatest enrichment of AP-1 motifs at ERα binding sites established at the highest dose of E2, which exhibit lower enrichment of EREs (10 nM; Fig. 3C; Fig. S4A). Comparison of published c-Fos ChIP-seq data with our ERα ChIP-seq data showed the greatest enrichment of c-Fos at ERα binding sites established with the highest doses of E2 (Fig. S4, B and C), suggesting that AP-1 actions at ERα enhancers, especially those lacking an ERE, may be a pharmacological response. Previous genomic studies found a strong negative correlation between ERE and AP-1 motifs (Carroll et al., 2006), also hinting at the types of observations that we have made here.

### Distinct RNA Pol II-dependent mechanisms at pM responsive and nM responsive promoters

We previously showed that E2-regulated genes, in aggregate, exhibit hormone-dependent loading of RNA polymerase II (Pol II) [i.e., preinitiation complex (PIC) formation], as well as promoter proximal Pol II pausing (Hah et al., 2011), which represent distinct mechanisms of gene regulation (Core and Adelman, 2019). To determine if promoters responsive to pM or nM concentrations of E2 might exhibit distinct Pol II-dependent mechanisms of gene regulation, we examined PIC formation and Pol II pausing by PRO-seq at representative examples of pM responsive (e.g., *ADRB1*) and nM responsive (e.g., *PGR*) genes. These genes showed clear and distinct Pol II dynamics, with *ADRB1* exhibiting paused Pol II in the absence of E2, transitioning into elongating Pol II with 10 pM or 10 nM of E2 (Fig. 6A, *left*). In contrast, *PGR* exhibited an absence of Pol II in the absence of E2, with loading and pausing of Pol II at 10 pM of E2, and additional pausing and a transition into elongating Pol II at 10 nM of E2 (Fig. 6A, *right*). These same trends were observed on a global scale, looking across all genes in each dosage group (Fig. 6B). The pM responsive genes showed a greater accumulation of reads in the promoter region (TSS to +500 pb) prior to E2 treatment, as well as a higher pausing index (reads in the paused regions/gene body reads) (Fig. 6C). Moreover, the pM responsive genes showed a decrease in the pausing index with the lowest doses of E2, whereas the nM responsive genes showed an increase in the pausing index at higher doses (Fig. 6D).

**Figure 6.**
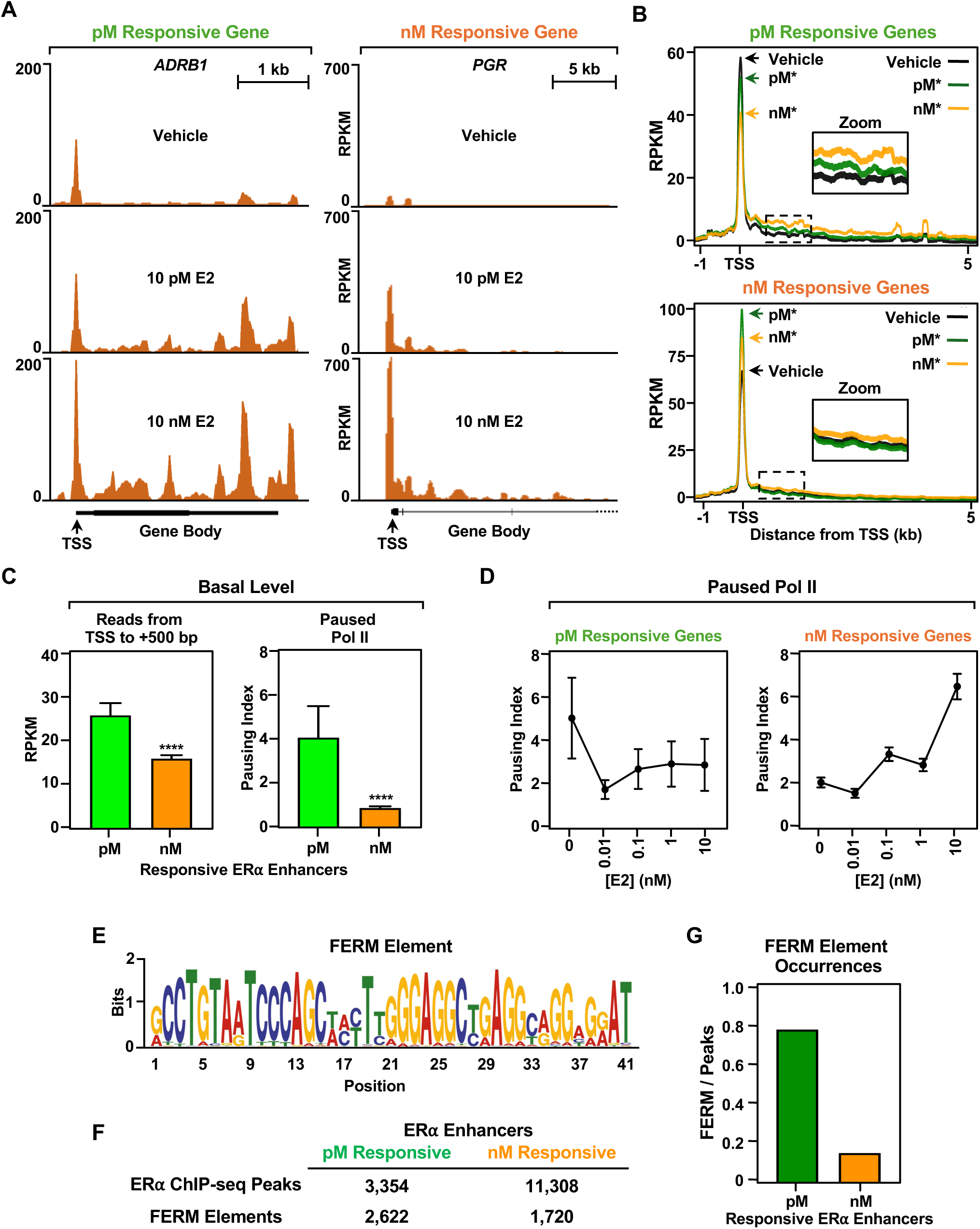
RNA Pol II pausing and PIC formation at the promoters of pM responsive and nM responsive genes. **(A)** Browser tracks showing representative genes in the pM (*ADRB1*) and nM (*PGR*) responsive gene groups. Transcription start sites (TSS, arrows) and gene bodies are indicated for each gene. **(B)** Metaplots showing the average profile of PRO-seq data near the TSS and first 5 kb of the gene body for pM (*top*) and nM (*bottom*) E2 dosage groups. *Insets:* enlarged images of the regions indicated by dashed box are shown to highlight transcription in the gene body. **(C)** Bar graphs showing read counts for TSS to +500 bp (*left*) and the pausing index (read counts at TSS/read counts in gene body; *right*) shown for pM responsive and nM responsive genes prior to E2 treatment. Each bar represents the mean + SEM. Asterisks indicate significant differences; Wilcoxon Rank Sum test, **** p < 5 × 10^-15^. **(D)** Line graphs showing pause indexes across a range of E2 treatment concentrations for pM responsive (*left*) and nM responsive (*right*) genes. Each point represents the mean ± SEM; n = 1,573 genes for pM and n = 2,352 genes for nM. **(E)** Motif logo for the human FERM element. **(F)** Table showing the number of ERα peaks (ChIP-seq) and FERM element motif occurrences (FIMO) shown for pM responsive and nM responsive ERBS. **(G)** Bar graph showing the number of occurrences of FERM elements per number of ERα peaks shown for pM responsive and nM responsive ERBS.

Integration of genomic data from pM responsive and nM responsive ERα enhancers and their corresponding target gene promoters reveals a clear dose shift for the genomic features assayed herein (Fig. S5). It also showed that enhancer transcription, measured by PRO-seq, exhibits similar dose sensitivity as target gene transcription, again with a rightward dose shift for the nM responsive genes (Fig. S5). Following these analyses, we explored additional aspects of enhancer transcription and enhancer RNA (eRNA) production in more detail. We previously showed that some eRNAs produced from ERα enhancers are enriched with functional eRNA regulatory motifs (FERMs) (Fig. 6E) that bind coregulators and help establish transcriptionally active enhancers (Hou and Kraus, 2022). We observed that FERMs are dramatically enriched in the eRNAs produced from enhancers established in response to pM levels of E2 compared to eRNAs produced from enhancers established in response to nM levels of E2 (Fig. 6F), especially when expressed as FERMs per ERα binding site (Fig. 6G). Together with the data from Figs. 1 and 2, these results show that the genes most highly responsive to low levels of E2 are regulated by ERα enhancers that form in open regions of chromatin and produce eRNAs enriched in FERMs, and have promoters with low-dose responsive paused Pol II.

### Differential coregulator association with chromatin-bound ER**α** at pM and nM doses of E2

To assess broadly how coregulator recruitment to chromatin-bound ERα is affected by different doses of E2, we mined our ERα RIME data from experiments conducted at low (10 pM) and high (10 nM) concentrations of E2. We had good (58%) coverage of ERα peptides by mass spectrometry in the RIME assays, indicating the protocol was working well (Fig. S6A). As noted above, we observed E2-dependent recruitment of ERα to chromatin, with similar enrichment at both 10 pM and 10 nM E2 (Fig. 4D, *top*). We identified hundreds of proteins associating with chromatin-bound ERα, some with maximal association at 10 pM E2 (Fig. S6B) and others with maximal association at 10 nM E2 (Fig. S6C). Both sets of proteins were enriched for GO terms related to mRNA processing, splicing, and nucleosome assembly (Fig. S6, D and E). The classical ERα coregulators p300 and NCOA3/SRC3 showed maximal (saturated) association with chromatin-bound ERα at 10 pM E2 (Fig. S6F), while NCOA2/SRC2 showed maximal (saturated) association with chromatin-bound ERα at 10 nM E2 (Fig. S6F). Comparison of our ERα RIME data with our previous mass spectrometry data for FERM-interacting proteins (Hou and Kraus, 2022) identified a set of proteins, many of which are involved in RNA processing and splicing, that may be associated with ERα enhancers in an E2 dose-dependent manner (Fig. S7).

### Poor outcomes of breast cancer patients on aromatase inhibitor treatment is associated with pM responsive ER**α** enhancers

To determine if E2 dose-dependent gene expression and enhancer formation is associated with clinical outcomes, we compared our genomic data from MCF-7 cells with gene expression and ERα cistromic data obtained from breast cancer patients. Overall survival during a 5-year period was significantly better in patients with higher expression of the nM responsive gene set, but not the pM responsive gene set (Fig. 7A). Focusing on expression of the mRNAs encoding FOXA1, GATA3, and ERα (*ESR1*), we observed that higher expression of *FOXA1*, but not *GATA3,* was associated significantly with a poorer outcome in patients with luminal A/B breast cancers (Fig. 7B). When examined together, elevated expression of *FOXA1* and *GATA3* were only associated significantly with a poorer outcome in patients expressing *ESR1* (Fig. 7B), consistent with the role of FOXA1 and GATA3 acting as pioneer factors to support ERα binding.

**Figure 7.**
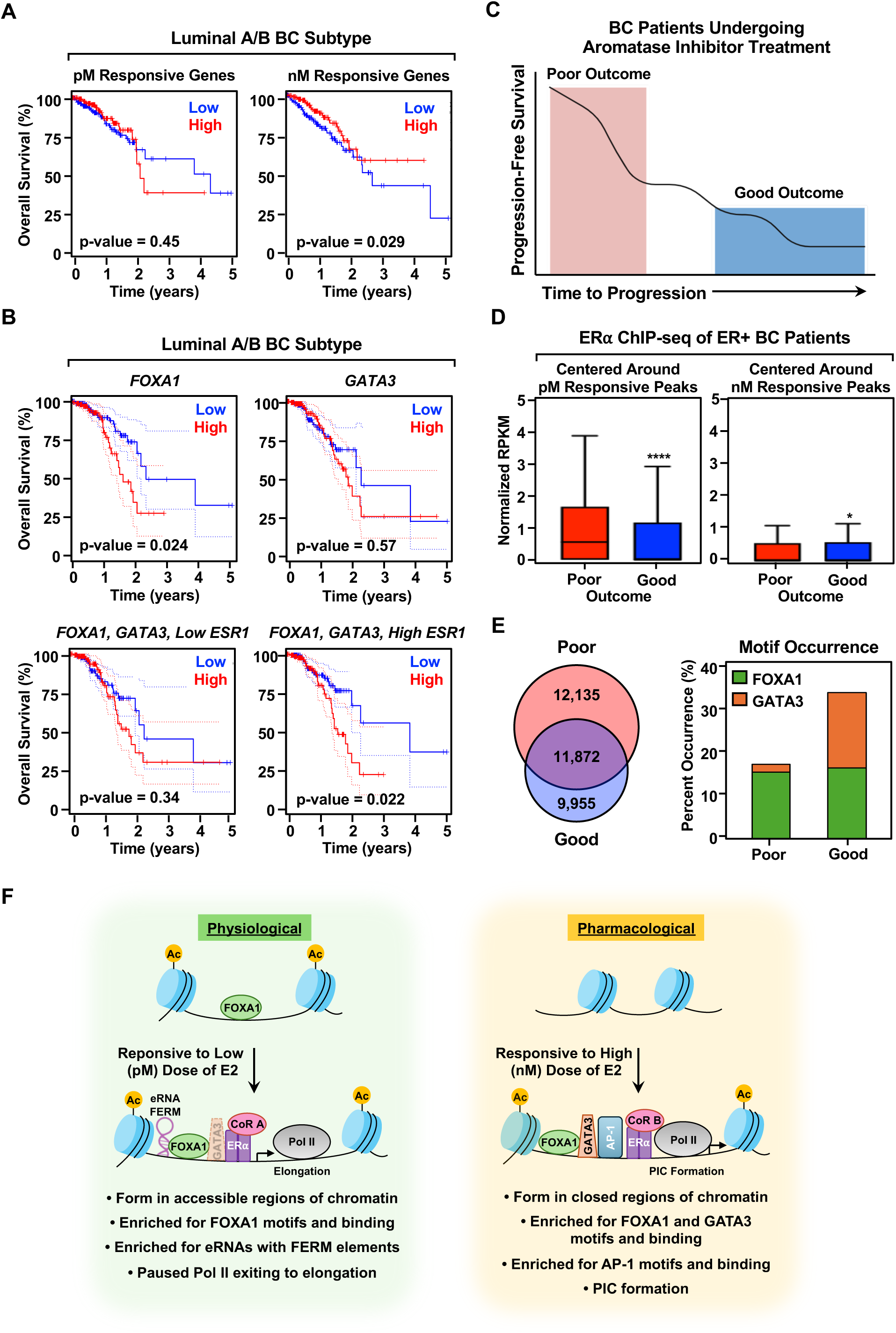
Outcomes of breast cancer patients on aromatase inhibitor treatment is associated with ERα binding at pM responsive enhancers. **(A)** Kaplan-Meier plots showing overall survival of breast cancer patients with high or low expression of the top 30 genes within the pM responsive (*left*) or nM responsive (*right*) gene sets based on RNA-seq. **(B)** Kaplan-Meier plots showing overall survival of patients with luminal A/B breast cancers with high or low expression levels of *FOXA1*, *GATA3*, *ESR1* mRNAs, alone or in combination as indicated. **(C)** Schematic diagram showing outcomes in patients who underwent aromatase inhibitor treatment: Poor outcome (time to progression < 12 months; highlighted in red) or good outcome (time to progression >24 months; highlighted in blue). Adapted from (Jansen et al., 2013). **(D)** Boxplots showing ERα binding by ChIP-seq within pM responsive (*left*) or nM responsive (*right*) ERBS for breast cancer patients with poor or good outcomes after aromatase inhibitor treatment. ChIP-seq data from breast cancer patients (Jansen et al., 2013) was mapped to the pM responsive or nM responsive ERBS determined in MCF-7 cells. Asterisks indicate significant differences; Wilcoxon Rank Sum test, * p < 5 × 10^-5^, **** p < 5 × 10^-15^. **(E)** Venn diagram showing the number of overlapping peaks between patient outcome groups (*left*) and a graph showing the occurrence of FOXA1 or GATA3 motifs near the ERα enhancers in breast cancers from patients with poor or good outcomes (*right*). **(F)** Model showing the dose-dependent effects of E2 on gene regulation by ERα in breast cancer cells. See the text for details.

Next, we investigated ERα enhancers and outcomes in breast cancer patients who underwent therapeutic suppression of circulating E2 via aromatase inhibitors by mining a publicly available data set (Fig. 7C) (Jansen et al., 2013). In this analysis, we assessed ERα binding at the pM responsive and nM responsive enhancers that we mapped in MCF-7 cells using ChIP-seq data from the breast cancer patients. Interestingly, we found that patients with a poor outcome had significantly higher ERα binding at pM responsive sites compared to patients with a good outcome (Fig. 7D). In contrast, patients with a poor outcome had a modest, but significant, reduction in ERα binding at nM responsive sites compared to patients with a good outcome (Fig. 7D). Likewise, we found that patients with a poor outcome had significantly higher enrichment of FOXA1 motifs versus GATA3 motifs near the ERα binding sites compared to patients with a good outcome (Fig. 7E). Together these results suggest that FOXA1-dependent, low dose-sensitive ERα enhancers respond to the circulating E2 that remains in breast cancer patients who do not respond well to aromatase inhibitor treatment. The pM sensitive ERα enhancers can drive gene expression programs that facilitate breast cancer growth and proliferation.

## Discussion

In this study, we explored a largely overlooked aspect of steroid hormone signaling, namely the dose-dependent effects of the hormone, especially in the context of the genomic actions of nuclear steroid hormone receptors. By exploring the molecular, genomic, and sequence features of ERα enhancers established over six orders of magnitude of E2 treatment, we were able to identify and characterize a class of ERα enhancers that are established and are active at low, physiological doses of E2 (1-100 pM). These ‘low dose’ enhancers are distinct from ‘high dose’ ERα enhancers established exclusively in the presence of high, pharmacological doses of E2 (100 nM) with respect to the molecular mechanisms, target gene sets, and biological outcomes. The implications of these findings, which are discussed in detail below, are manifold.

### Mechanisms of action and biological consequences of low dose estrogen signaling

For most of a woman’s adult life, exposure to E2 typically occurs at circulating levels in the pM range (typically ∼100 pM up to 2 nM, but frequently lower in prepubertal or postmenopausal females). Most of what we know, however, about the genomic actions of E2 are from cell-based studies conducted at 100 nM of E2. This has left a significant gap in knowledge about estrogen signaling and more broadly about steroid hormone signaling. Our studies have defined the molecular features of pM responsive and nM responsive ERα enhancers, and highlighted key differences between the two (Fig. 7F). ERα in the presence of low, physiological concentrations of E2 binds preferentially to EREs located in open and accessible regions of chromatin prepared by FOXA1. Activation of these enhancers leads to the production of eRNAs containing FERM elements, as well as the subsequent activation of target gene promoters loaded with paused RNA Pol II. These genomic features are hallmarks of the most permissive regulatory element-gene pairs. Thus, physiological E2 signaling supports a gene regulation program with the lowest bar for activation. In contrast, high dose, pharmacological E2 signaling requires GATA3 to open new, less tractable regions of chromatin, where ERα promotes the loading of RNA Pol II through PIC formation (Fig. 7F).

Our analyses suggest that these distinctions can have important biological outcomes. Cell growth and GO analyses indicate that different doses of E2 control biologically distinct outcomes, with low doses of E2 very effective in promoting proliferation in breast cancer cells (Fig. 1, D and E). As such, physiological, pathological, or therapeutic conditions with low circulating levels of E2 may experience biology-altering, hormone-dependent gene expression outcomes. For example, breast cancer patients with poor outcomes on aromatase inhibitor therapy may experience incomplete reductions in circulating E2. As our analyses suggest, the breast cancers in these patients may capitalize on the remaining E2 levels to promote enhancer formation at low dose-responsive ERα binding sites leading to enhanced cancer cell growth (Fig. 7, C through E). Our results bring attention to the important biological outcomes that can be driven by low, subsaturating levels of hormone.

### Considerations for the study of steroid hormone signaling

The advent of genomics over two decades ago led to the discovery of new facets of steroid hormone signaling and the molecular mechanisms of nuclear receptor function. ERα was the first nuclear receptor to be subject to genome-wide localization studies (Carroll et al., 2005; Carroll et al., 2006), with a set of standard conditions established early on and applied fairly consistently since then. A key parameter that has been used seemingly invariably in such studies is a high dose (typically 100 nM) of hormone. With an affinity (Kd) of ERα for E2 of ∼0.1 nM (Kuiper et al., 1997; Lin et al., 2013), the fractional occupancy of ERα at 10 to 100 pM is ∼10 to 50 percent. As our results show, these subsaturating levels of E2 are sufficient to drive the biology of the ERα and do so in a fundamentally different manner than high, supersaturating doses of E2. Consequently, previous studies conducted at high E2 concentrations captured combined effects, preventing clear differentiation of mechanisms occurring at different doses. In this regard, previous studies may have overemphasized the importance of GATA3 and AP-1 in physiological responses to E2, although both are likely to play important roles in pharmacological and therapeutic responses to E2.

A recent study examining the dose-dependent effects of androgens in androgen receptor (AR)-expressing cells reported observations different than ours with E2/ERα, likely reflecting differences between the cell biology of AR and ERα. Safi *et al*. observed no significant binding of AR to chromatin by ChIP-seq at low doses of the synthetic androgen R1881 (10 and 30 pM) that elicited proliferative responses (Safi et al., 2024). This is in contrast to the significant binding of ERα to chromatin that we observed at 1 and 10 pM of E2. Based on additional analyses, they concluded that low doses of androgens act through an AR monomer to promote non-genomic activation of the mTOR signaling pathway to drive proliferation. Only at higher doses do androgens promote the classical genomic effects of AR to drive an androgen-dependent transcriptional program (Safi et al., 2024). An important distinction between the E2/ERα and androgen/AR systems that is likely to underlie the differences at low doses of hormone is that ERα is nuclear in the absence of ligand, whereas AR is cytosolic and must translocate to the nucleus upon hormone exposure. These results highlight the need to evaluate steroid hormone signaling over a range of doses and explore a variety of potential molecular mechanisms.

### Potential role of low dose-responsive ER**α**enhancers in the evolution of the estrogen response

Our work highlights the convergence of retrotransposons, regulatory elements, open chromatin, and transcriptional enhancers, with implications for the evolution of a signal-regulated transcriptional response. Many transcription factor binding sites, including those for ERα, are located in Alu and related elements (Norris et al., 1995; Polak and Domany, 2006; Su et al., 2014). In a previous study, we discovered the eRNA FERM element, which is encoded in Alu elements (humans) and B1 elements (rodents and other mammals), two types of retrotransposons belonging to the class of short interspersed nuclear elements (SINEs) (Hou and Kraus, 2022). Isolated FERM RNAs are sufficient to drive E2-regulated target gene expression when artificially recruited to and tethered at native ERα enhancers (Hou and Kraus, 2022). The combination of EREs and FERMs – two sequences that are hallmarks of low dose-responsive ERα enhancers – within Alu elements comprises a genetic element with the potential to mobilize and populate new regions of the human genome. This could allow for reorganization of the low dose estrogen responsive ERα cistrome and transcriptome, ultimately generating variability in the hormone response across the human population. Since evolution is most likely to act in the context of physiological, rather than pharmacological, doses of E2, low dose-responsive ERα enhancers have the potential to shape the natural history and evolution of the estrogen response, as well as therapeutic responses that target the synthesis of E2 or the activity of ERα.

### ‘Dose’ as a driver of enhancer function and transcriptional regulation

Many aspects of transcriptional regulation are quantitatively encoded rather than all or none, with cells capable of sensing and interpreting the concentration or abundance of signaling molecules and effectors. Our studies on the effects of hormone dose in ERα-dependent transcriptional regulation provide a clear example of this. More broadly across the realm of enhancer-mediated gene regulation, the quantitative abundance of inputs such as ligands, transcription factors, and coregulators have the potential to elicit dose-dependent effects that control gene expression outputs. For example, a recent study has shown that some regulatory elements are sensitive to the dosage of the transcription factors that control them (Naqvi et al., 2023). The results of another study suggest that transcription factors compete for limiting amounts of coregulators, such as CREB binding protein (CBP), which exhibits haploinsufficiency (Kamei et al., 1996), akin to a dose-dependent effect. As these types of analyses evolve, concepts related to dose sensitivity may be found to extend to post-translational modifications, eRNAs, and nucleic acid-based motifs.

## Experimental Procedures

### Antibodies

The custom rabbit polyclonal antiserum raised against the first 113 amino acids of human ERα was generated in-house as previously described (Kraus and Kadonaga, 1998). Other antibodies used were as follows: FOXA1 (Abcam, ab23738; RRID:AB_2104842), GATA3 (Abcam, ab199428; RRID:AB_2819013), histone H3K27ac (Active Motif, 39133; RRID:AB_2561016), and rabbit IgG isotype control (Thermo Fisher Scientific, 10500C; RRID:AB_2532981).

### Cell culture and treatments

MCF-7 cells (RRID:CVCL_0031) were kindly provided by Dr. Benita Katzenellenbogen (University of Illinois, Urbana-Champaign, Champaign, IL) and were maintained in minimal essential medium (Sigma-Aldrich, M1018) supplemented with 5% calf serum (Sigma, C8056), 100 U/mL penicillin-streptomycin (Gibco, 15140122), and 25 μg/mL gentamicin (Gibco, 1571004). MCF-7 cells were authenticated for cell-type identity using the GenePrint 24 system (Promega, B1870), blotting for ERα, and confirmed as Mycoplasma-free every 6 months using the Universal Mycoplasma Detection Kit (ATCC, 30-1012K). Fresh cell stocks were regularly replenished from original stocks every few months (no more than 10 passages). For experiments involving estrogen treatment, the cells were grown for 3 days in phenol red-free Minimum Essential Medium Eagle (Sigma, M3024) supplemented with 5% charcoal-dextran-treated calf serum (CDCS; Sigma, C8056), 1 mM GlutaMAX (Gibco, 35050061), 100 U/mL penicillin-streptomycin (Gibco, 15140122), and 25 μg/mL gentamicin (Gibco, 1571004). All subsequent experiments involving E2 treatment were carried out in estrogen-free medium. For induction experiments, cells were treated with the indicated concentration of E2 (0 to 100 nM; Sigma, E8875) or DMSO vehicle for the indicated experimental time.

### Cell growth assays

MCF-7 cells were plated in phenol red-free MEM containing 5% CDCS for 72 hours prior to growth assays, as previously described (Kim and Kraus, 2025). Cells were plated at a density of 20,000 cells per well in a 6-well dish and treated with the indicated concentration of E2 for 5 days. Medium containing E2 was replenished every 2 days. At the experiment endpoint, the cells were trypsinized and live cell count was obtained by staining with 0.4% trypan blue (Sigma, T8154) and counting using the BioRad TC20 automated cell counter.

### siRNA-mediated knockdown

siRNA-mediated knockdown was performed as previously described (Kim and Kraus, 2025) using an SE Cell Line 4D-Nucleofactor X kit (Lonza; V4XC-1032; EEID:SCR_023155) according to the manufacturer’s protocol. Briefly, MCF-7 cells were plated in phenol red-free MEM containing 5% CDCS. siRNAs targeting the mRNAs of interest or a control siRNA (Select Negative Control No. 1; Thermo Fisher Scientific, 4390843) were applied to MCF-7 cells at a final concentration of 30 nM in SE transfection reagent before electroporation using a Lonza 4D Nucleofector Nucleofection Transfection System Core Unit & X Unit with the CA 137 program. Cells were used for various assays 48 hours after siRNA transfection.

#### siRNA sequences

We used the following nucleic acid oligonucleotides for targeted knockdowns:

siFOXA1_Hs01_00168403 5’-CACACAAACCAAACCGUCA [dT][dT]-3’
siFOXA1_ Hs01_00168403_AS 5’-UGACGGUUUGGUUUGUGUG [dT][dT]-3’
siGATA3_Hs01_00153931 5’-CUCUGGAGGAGGAAUGCCA [dT][dT]-3’
siGATA3_Hs01_00153931_AS 5’-UGGCAUUCCUCCUCCAGAG [dT][dT]-3’

### Preparation of cell extracts and Western blotting

#### Preparation of whole cell lysates

At treatment endpoints, the cells were washed twice with ice-cold 1x PBS, scraped, and pelleted. The cell pellets were lysed in Lysis Buffer [20 mM Tris-HCl pH 7.6, 150 mM NaCl, 1 mM EDTA, 1% NP-40, 1% sodium deoxycholate, 0.1% SDS, 1 mM DTT, 1x protease inhibitor cocktail (Roche, 11697498001)] by vortexing for >30 seconds. The lysate was cleared of debris by centrifugation at >20,000 x g for 15 minutes at 4°C. The supernatant containing whole-cell extract was used for further analyses.

#### Subcellular fractionation

At treatment endpoints, the cells were washed twice with ice-cold 1x PBS, scraped, and pelleted. The cells were incubated with 100x packed cell volume (PCV) of Lysis Buffer (10 mM Tris pH 7.5, 60 mM KCl, 15 mM NaCl, 0.34 M sucrose, 2 mM EDTA, 0.5 mM EGTA, 0.1% Triton X-100, 650 μM spermidine, 1 mM DTT, 1x protease inhibitor cocktail) for 10 minutes on ice. The cells were collected at 400 x *g* for 5 minutes, and the supernatant was collected as the cytoplasmic fraction. Pelleted nuclei were washed twice in excess Wash Buffer (10 mM Tris pH 7.5, 10 mM KCl, 0.34 M sucrose, 1.5 mM MgCl_2_, 1 mM DTT, 1x protease inhibitor cocktail). Pelleted nuclei were resuspended in 20x PCV of Nuclear Lysis Buffer (50 mM Tris pH 7.5, 140 mM NaCl, 1.5 mM MgCl_2_, 0.5% NP-40, 1 mM DTT, 1x protease inhibitor cocktail) for 20 minutes on ice, with occasional vortexing. Nuclear lysates were then cleared at >20,000 x *g* for 10 minutes at 4°C. The resulting supernatant was collected as the nucleoplasmic fraction. Pelleted chromatin was washed twice in excess Chromatin Buffer (10 mM HEPES pH 8, 2 mM MgCl_2_, 1% SDS, 1 mM DTT, 1x protease inhibitor cocktail). The chromatin was then pelleted and resuspended in 2x PCV of Chromatin Buffer supplemented with 1 U/μL of universal nuclease (ThermoFisher, 88700). The resuspended material was then incubated for 1 hour at 37°C with shaking at 750 rpm in a thermomixer. The resulting material was used as the chromatin fraction.

#### Determination of protein concentrations and Western blotting

Protein concentrations were determined using a BCA protein assay (Pierce, 23225). Aliquots of extract for analysis by SDS-PAGE were mixed with 4x SDS Loading Solution (250 mM Tris pH 6.8, 40% glycerol, 0.04% Bromophenol Blue, 4% SDS, 1% β-mercaptoethanol) to a final concentration of 1x. The samples were heated to 95°C for 5 minutes before separating by SDS-PAGE and transferring to nitrocellulose membranes. The membranes were blocked using 5% non-fat milk in TBST for 30 minutes, incubated with the indicated primary antibody in TBST overnight at 4°C, followed by incubation with goat anti-rabbit HRP-conjugated IgG (1:5000 dilution). The washed blots were treated with ECL detection reagent (Thermo Fisher Scientific, 34077, 34095) and visualized using a BioRad ChemiDoc system at multiple exposure times.

### Reverse transcription-quantitative PCR (RT-qPCR)

MCF-7 cells were subjected to E2 dose treatments as described above and collected at the indicated experimental timepoint. For the reverse transcription quantitative PCR (RT-qPCR) assays, cells were treated with E2 at the indicated dose for 2 hours. Total RNA was isolated using the QIAGEN RNeasy Plus Mini Kit (Qiagen, 74136) according to the manufacturer’s protocol. Total RNA was reverse transcribed using oligo(dT) primers and MMLV reverse transcriptase (Promega, M1705) to generate cDNA. The resulting cDNA pool was treated with 3 units of RNase H (Enzymatics, Y9220L) for 30 minutes at 37°C, and then analyzed by qPCR using a Roche LightCycler 480 system with SYBR Green detection and gene-specific primers (see list below). Target gene expression was normalized to the expression of GAPDH mRNA.

#### RT-qPCR primer sequences

We used the following nucleic acid oligonucleotides for qPCR:

GAPDH-F 5’-GTCTCCTCTGACTTCAACAGCG-3’
GAPDH-R 5’-ACCACCCTGTTGCTGTAGCCAA-3’
PGR-F 5’-ACCACGCACGTTCTGCTAAT-3’
PGR-R 5’-TGGAGCACCTAAGAGGAGCA-3’
ADRB1-F 5’-TTCCTGCCCATCCTCATGCACT-3’
ADRB1-R 5’-GTAGAAGGAGACTACGGACGAG-3’
WNT16-F 5’-TCGGAAACACCACGGGCAAAGA-3’
WNT16-R 5’-GCGGCAGTCTACTGACATCAAC-3’
MBOAT1-F 5’-GGTTTCCACAGCTTGCCAGAAC-3’
MBOAT1-R 5’-ACCAGTCATCCACAAGGCAGGT-3’
TSKU-F 5’-AGTCGCTTGACCTCAGCCACAA-3’
TSKU-R 5’-TCGTGAAGGCAGACACTGAGAC-3’
GREB1-F 5’-GGTCTGCCTTGCATCCTGATCT-3’
GREB1-R 5’-TCCTGCTCCAAGGCTGTTCTCA-3’
CYP1B1-F 5’-GCCACTATCACTGACATCTTCGG-3’
CYP1B1-R 5’-CACGACCTGATCCAATTCTGCC-3’
MIDEAS-F 5’-GCACAAGCCATCAGTCATCGTC-3’
MIDEAS-R 5’-CTGCTTTGGTTTCCGCACGGAA-3’
TFAP2C-F 5’-CACCTGTTGCTGCACGATCAGA-3’
TFAP2C-R 5’-AGGAGCGACAATCTTCCAGGGA-3’

### Sequencing data quality control

For all of the NextGen sequencing assays described below (RNA-seq, ChIP-seq, ATAC-seq, and PRO-seq), raw FASTQ files of the data were assessed for sequencing quality using FastQC (Andrews, 2010). Quality metrics, such as total reads, correlation between replicates, per-base sequence quality, GC content, adapter content, and duplication levels were evaluated across all samples to ensure data suitability for downstream analysis per the Encyclopedia of DNA Elements (ENCODE) Project standards (Consortium et al., 2020).

### RNA-sequencing and analysis

#### Generation of RNA-seq libraries and sequencing

At treatment endpoints, RNA-seq libraries were generated from two biological replicates per condition as previously described (Hou and Kraus, 2022). Total RNA from MCF-7 cells treated as indicated was isolated using the Qiagen RNeasy Plus Mini kit (Qiagen, 74136) according to the manufacturer’s protocol. mRNA was enriched using Oligo(dT)25 beads (Invitrogen, 61002) and strand-specific, single-end libraries were generated using unique Illumina barcodes with a final amplification of 13 cycles. Resulting libraries were subjected to QC analyses, including assessment of yield by Qubit fluorometer (Invitrogen, Q32851) and size distribution by DNA ScreenTape system (Agilent Technologies, G2964AA, 5067-5584, and 5067-5603), and sequenced using an Illumina NextSeq 2000 by 100 bp single-end sequencing.

#### Analysis of RNA-seq data

FASTQ files were subjected to quality control analyses using the FastQC tool (Andrews, 2010). The single-end RNA-seq reads were aligned to the human genome using TopHat v2.0.12 (Kim et al., 2013), with genome indices built using Bowtie2 v2.1.0 (Langmead et al., 2009). Aligned bam files were used for downstream processing (Trapnell et al., 2009). Aligned reads were quantified using featureCounts from the Subread v1.6.3 package (Liao et al., 2014). Gene-level read counts were obtained using the GENCODE v35 GTF annotation, specifying exon-based counting (-t exon), strand specificity (-s 2), and fractional assignment of multi-mapped reads (--fraction). Gene counts were normalized and tested for differential expression using DESeq2 (Love et al., 2014). Normalized counts were used for principal component analysis (PCA) and hierarchical clustering. Pairwise comparisons between vehicle and each E2 dose were performed using the Wald test in DESeq2. Differentially expressed genes (DEGs) were defined by adjusted p-value <0.05 and a fold change >1.5 (upregulated) or <0.67 (downregulated).

### Chromatin immunoprecipitation (ChIP) and ChIP-qPCR

#### Chromatin immunoprecipitation

Chromatin immunoprecipitation (ChIP) was performed as previously described (Hou and Kraus, 2022)with some modifications. MCF-7 cells were grown to ∼80% confluence under conditions and with E2 treatments as described above. After treatment, the cells were crosslinked with 1% formaldehyde (Thermo Fisher Scientific, 28-906) in 1x PBS at 37°C for 10 minutes. The crosslinking was quenched with glycine at a final concentration of 125 mM for 5 minutes at 4°C. The cells were washed thoroughly with ice-cold 1x PBS and collected by scraping into 1 mL of ice-cold 1X PBS. The cells were pelleted by centrifugation and lysed by pipetting in Farnham Lysis Buffer (5 mM PIPES pH 8, 85 mM KCl, 0.5% NP-40, 1 mM DTT, and 1x complete protease inhibitor cocktail). Nuclei were collected by brief centrifugation and resuspended in SDS Lysis Buffer (50 mM Tris-HCl pH 7.9, 1% SDS, 10 mM EDTA, 1 mM DTT, and 1x complete protease inhibitor cocktail) by pipetting and incubating on ice for 10 minutes.

The chromatin was sheared to ∼200 to 500 bp DNA fragments by sonication using a Diagenode Pico sonication device (Diagenode, B01080010) for 8 cycles of 30 seconds on/30 seconds off. Fragment size was verified by agarose gel electrophoresis before quantification of protein concentrations using a BCA protein assay kit (Pierce, 23225). The samples were diluted 10-fold using Dilution Buffer (20 mM Tris-HCl pH 7.9, 0.5% Triton X-100, 2 mM EDTA, and 150 mM NaCl) with freshly added 1 mM DTT and 1x protease inhibitor cocktail. Soluble chromatin (500 μg per reaction) was precleared with Protein A Dynabeads (Invitrogen, 10001D). Protein A Dynabeads were incubated with 10 μL of polyclonal antiserum for ERα or 5 μg of rabbit IgG control (Thermo Fisher Scientific, 10500C; RRID: AB_2532981) for 2 hours, washed with Dilution Buffer, and applied separately to precleared chromatin samples. The chromatin fractions and the antibody-loaded Protein A Dynabeads were incubated overnight at 4°C.

The immune complexes on the beads were washed once with each of the following Wash Buffers in order: (i) Low Salt Wash Buffer (20 mM Tris-HCl pH 7.9, 2 mM EDTA, 125 mM NaCl, 0.05% SDS, 1% Triton X-100, and 1x complete protease inhibitor cocktail); (ii) High Salt Wash Buffer (20 mM Tris-HCl pH 7.9, 2 mM EDTA, 500 mM NaCl, 0.05% SDS, 1% Triton X-100, and 1x complete protease inhibitor cocktail); (iii) LiCl Wash Buffer (20 mM Tris-HCl pH 7.9, 1 mM EDTA, 250 mM LiCl, 1% NP-40, 1% sodium deoxycholate, and 1x complete protease inhibitor cocktail); (iv) 1x Tris-EDTA (TE) containing 1x complete protease inhibitor cocktail. The immune complexes precipitates were transferred to a new tube in 1x TE before incubation in Elution Buffer (100 mM NaHCO_3_ and 1% SDS) overnight at 65°C to de-crosslink and elute. Eluates were treated with DNase-free RNase (Roche, 11119915001) for 30 minutes at 37°C, followed by 50 μg proteinase K (Life Technologies, 2542) for 2 hours at 55°C. Genomic DNA was purified by phenol chloroform extraction.

#### ChIP-qPCR

For ChIP-qPCR assays, MCF-7 cells were treated with the indicated concentration of E2 for 30 minutes. ChIPed DNA was analyzed by qPCR as described for RT-qPCR above. Nonspecific background signals were determined by performing ChIP using rabbit IgG. The data were expressed as a percent of input or a relative enrichment versus control or background.

#### ChIP-qPCR primer sequences

We used the following nucleic acid oligonucleotides for ChIP-qPCR:

PGR-enhancer-F: 5’-CACTTGTCACATTGGAGGCAG-3’
PGR-enhancer -R: 5’-TGGGTCATCCTGTTCCCTTTG-3’
ADRB1-enhancer -F: 5’-AGGGCTGGTAGTTAGGAGGA-3’
ADRB1-enhancer -R: 5’-AGCTTTTCTATGGGAGATGTGCT-3’
WNT16-enhancer -F: 5’-GCTTTCCCCCAAGGTTGCTA-3’
WNT16-enhancer -R: 5’-CTCGGGACCTCAGGCTCTTA-3’
MBOAT1-enhancer -F: 5’-CTTCCAAACTCGCAAGCCAC-3’
MBOAT1-enhancer -R: 5’-CTCCAGCAGGAGTGAGTGTG-3’
TSKU-enhancer -F: 5’-TTCCCCGGAGTTTACTTGCC-3’
TSKU-enhancer -R: 5’-AAAGTCCTCTGACCCCCTGT-3’
GREB1-enhancer -F: 5’-TAGGCTTCAAGAGGACCACA-3’
GREB1-enhancer -R: 5’-AGCAGCAAAACTGCATAGGA-3’
CYP1B1-enhancer -F: 5’-CCGGGTTTTAAGGACTGGGT-3’
CYP1B1-enhancer -R: 5’-TGAAAGCCTGCTGGTAGAGC-3’
MIDEAS-enhancer -F: 5’-CCCAGAAGATCTGAGGGCAA-3’
MIDEAS-enhancer -R: 5’-ACATCCTGCCCAGTTTCCTC-3’
TFAP2C-enhancer -F: 5’-CCGTGACCCCGATTTTGGAT-3’
TFAP2C-enhancer -R: 5’-CCAGCTCACCTCGCAGTC-3’

### Chromatin immunoprecipitation (ChIP)-sequencing and analysis

#### Generation of ChIP-seq libraries and sequencing

ChIP-seq libraries were generated from two biological replicates for each condition. A total of 100 ng of ChIPed DNA, prepared as described above, were used for library generation, as described previously with few modifications (Hou and Kraus, 2022). Sequencing adaptors were annealed as previously described (Hou and Kraus, 2022) and the libraries were sequenced using an Illumina NextSeq 2000. At least two biological replicates were sequenced for each condition with a minimum of ∼40 million raw reads per condition.

#### Analysis of ChIP-seq data

Raw ChIP-seq reads were adapter-trimmed using cutadapt v1.9.1 (Martin, 2011) and aligned to the hg38 genome with Bowtie v1.0.0 (Langmead et al., 2009), allowing up to 2 mismatches and retaining uniquely mapped reads. Low-quality (MAPQ < 30), secondary, and unmapped reads were removed with samtools (Li et al., 2009), and duplicates were marked using Picard MarkDuplicates version 1.127 (http://broadinstitute.github.io/picard/). Strand-specific normalized coverage tracks were generated in R using the groHMM package (Chae et al., 2015), binned at 25 bp. UCSC-compatible wiggle files were converted to BigWig format using wigToBigWig for genome browser visualization (Kent et al., 2010). ChIP-seq peaks were identified using MACS v1.4.2 (Zhang et al., 2008) with default parameters. Signal enrichment around peak summits was visualized using deepTools (Ramirez et al., 2016; Zhang et al., 2008). The computeMatrix tool was used in reference-point mode (--referencePoint center) to calculate normalized coverage ±5 kb around MACS peak summits. The resulting matrix was visualized using plotProfile.

### Assay for transposase-accessible chromatin (ATAC)-sequencing and analysis

#### Generation of ATAC-seq libraries and sequencing

ATAC-seq libraries were generated as previously described (Huang et al., 2022) with a few modifications. MCF-7 cells were grown and treated as described above, collected by trypsinization at the experimental endpoint, and counted. Fifty thousand live cells were used for each experimental condition. Nuclei were isolated by resuspending the cells in Hypotonic Lysis Buffer (10 mM Tris pH 7.6, 10 mM NaCl, 3 mM MgCl_2_, 0.1% NP-40, 0.1% Tween-20, 0.001% digitonin) and incubating on ice for 3 minutes. The nuclei were then washed once with Hypotonic Lysis Buffer without NP-40 or digitonin. The nuclei were then resuspended in Transposition Mix (10 mM Tris pH 7.6, 5 mM MgCl_2_, 10% DMF, 100 nM Tn5 transposase, 50 mM NaCl, 1 mM KCl, 5 mM Na_2_HPO_4_, 20% Tween-20, 1% digitonin) and incubated at 37°C for 30 minutes with mixing at 1000 rpm. The resulting tagmented DNA fragments were purified by column purification using a Qiagen PCR purification kit (Qiagen, 28104). The final amplification of the libraries was performed using the NEBNext High-Fidelity Master Mix (NEB, M0541S), and Nextera i5 and i7 primers for a total of 7 cycles. Excess adapters were removed from the libraries by using 1.3x volume of AMPure Purification beads (Beckman Coulter, A63881) and the libraries were sequenced on an Illumina NextSeq 2000 sequencer by paired-end sequencing.

#### Analysis of ATAC-seq data

The paired-end ATAC-seq reads were aligned to the human genome (hg38) using BWA-MEM v0.7.15 (Li, 2013) with the -M flag to mark secondary alignments. Low-quality alignments (MAPQ < 10) were removed using samtools (Li et al., 2009) and alignments were sorted and deduplicated using Picard MarkDuplicates (REMOVE_DUPLICATES=true) (Institute, 2017; Li and Durbin, 2009). The final BAM files were indexed for downstream analyses (Institute, 2017; Li, 2013). ATAC-seq peaks were identified using MACS2 v2.1.0 (Zhang et al., 2008)with paired-end BAM input (-f BAMPE). The resulting peaks were used for downstream accessibility and motif enrichment analyses (Feng et al., 2012). ATAC-seq signal tracks were generated in R using the WriteWiggleNorm function from the groHMM package (Chae et al., 2015), using bed files. Read coverage was binned in 25 bp windows and normalized to the average library size across all conditions. The resulting UCSC-style wiggle (.wig) files were converted to BigWig format using the UCSC wigToBigWig tool, for visualization in genome browsers (Chae et al., 2015; Kent et al., 2010).

ATAC-seq peaks called by MACS2 were formatted into SAF (Simplified Annotation Format) using R and annotated with ChIPseeker (Yu et al., 2015). Genomic coordinates were converted into GRanges objects, and peaks were annotated relative to transcription start sites (TSS ±1 kb) using TxDb.Hsapiens.UCSC.hg38.knownGene and org.Hs.eg.db annotation databases. The distribution of peaks across genomic features (e.g., promoter, exon, intron, intergenic) was visualized with pie charts using plotAnnoPie (Carlson, 2023a; b). To examine chromatin accessibility at regions uniquely enriched in the ChIP-seq analyses, peak coordinates were extracted and used as reference points. Using the groHMM R package (Chae et al., 2015), read coverage from the ATAC-seq data was summarized across ±5 kb windows (100 bp bins) around these peak centers via the MetaGene function (Chae et al., 2015). Read counts were normalized both by the number of peaks and total reads per library, scaled to a common expected read depth. Resulting signal profiles were visualized using base R plotting functions.

### Precision run-on (PRO)-sequencing and analysis

#### Nuclear run-on and RNA hydrolysis

Nuclear run on and library preparation was performed following the PRO-seq protocol (Judd et al., 2020; Mahat et al., 2016) using approximately 5 million MCF-7 nuclei per reaction with a few modifications. Nuclei in 50 μL of Storage Buffer were resuspended in 50 μL of 2x ROMM Buffer [10 mM Tris-HCl pH 8, 5 mM MgCl_2_, 300 mM KCl, 40 μM each of Biotin-11-ATP/CTP/GTP/UTP (PerkinElmer, NEL544001EA, NEL542001EA, NEL545001EA, NEL543001EA), 1% Sarkosyl, 1 mM DTT, and 20U of SUPERase-In RNase inhibitor] per reaction. Run-on reactions were incubated for 5 minutes at 37°C with gentle mixing. RNA was extracted by TRizol LS (Invitrogen, 10296028), followed by ethanol precipitation. The resulting RNA was then sheared by base hydrolysis with 0.2 N NaOH for 10 minutes on ice and quenched with 500 mM Tris-HCl pH 6.8. Sheared RNA was subjected to salt exchange chromatography using Bio-Spin P30 columns (BioRad, 7326250).

#### Enrichment of nascent RNA and preparation of PRO-seq libraries

Libraries were prepared using two biological replicates per condition. Sheared RNA was ligated to the VRA3 adaptor at the 3’ end using RNA ligase (NEB, M0204S) before enriching by binding to base-activated streptavidin C1 magnetic beads (Invitrogen, 65002). Enriched RNA was then subjected to on-bead 5’ hydroxyl repair using T4 PNK (NEB, M0201) in the presence of ATP, followed by 5’ decapping by RppH (NEB, M0356S), and 5’ adaptor ligation to the VRA5 adaptor using RNA ligase. Ligated RNA was then eluted from the streptavidin beads by TRizol (Invitrogen, 15596018), followed by ethanol precipitation. Single strand cDNA was generated by reverse transcription using Maxima H Minus RT enzyme (Thermo Fisher Scientific, EP0751). The resulting cDNA was then amplified using universal small RNA and indexed small RNA primers for a total of 13 cycles. Excess adaptors were removed using SPRI beads (Beckman Coulter, A63880). The quality of resulting libraries was assessed using the TapeStation 4150 Bioanalyzer system. Libraries were sequenced using Illumina NextSeq 2000 by 100 bp single-end sequencing.

#### Analysis of PRO-seq data

The PRO-seq data quality was assessed using FastQC software (1). The data were then processed using a customized implementation of the proseq2.0 pipeline (Danko et al., 2015). The reads were aligned to the human genome (hg38) using the BWA aligner (Li and Durbin, 2009), excluding random chromosomes via a curated chromosome size file. The pipeline was executed in single-end, strand-specific mode (-SE -P −4DREG) for each sample. Strand-specific coverage was computed using deepTools bamCoverage v2.3.5 with 50 bp binning and RPKM normalization (Danko et al., 2015; Ramirez et al., 2016). The resulting bedGraph files were converted to the BigWig format using bedGraphToBigWig (Kent et al., 2010), ensuring compatibility with the filtered hg38 chromosome sizes for accurate genome browser visualization.

### Genomic data visualization

Heatmaps were created using Java Treeview (Saldanha, 2004) to visually display expression or enrichment that was significantly altered by the experimental conditions. Biovenn (Hulsen et al., 2008) was used to construct Venn diagrams for up- and downregulated genes or peaks, with overlaps. Boxplots were generated using custom R scripts to express FPKM values across the experimental conditions. Statistical significance among these comparisons was determined using Wilcoxon rank sum tests (p < 0.05).

### Rapid Immunoprecipitation Mass spectrometry of Endogenous proteins (RIME)

#### Immunoprecipitation

Rapid immunoprecipitation followed by mass spectrometry analysis was performed using MCF-7 cells as previously described (Papachristou et al., 2018) with a few modifications. Each condition had a minimum of four independent biological replicates. Approximately 10^8^ cells were used per experimental condition. Briefly, magnetic protein A Dynabeads were first blocked in 5 mg/mL BSA in 1x PBS for 30 minutes at room temperature. Polyclonal ERα antibody was added to the blocked beads overnight at 4°C with gentle mixing. The beads were then washed twice and resuspended in RIPA Buffer [50 mM HEPES pH 7.6, 1 mM EDTA, 0.7% (w/v) sodium deoxycholate, 1% (v/v) NP-40, 500 mM LiCl, 1x protease inhibitor cocktail] until further use.

At the end of the experiments, the cells were crosslinked using 1% formaldehyde for 10 minutes at room temperature and quenched adding glycine to a final concentration of 100 mM. The cells were washed twice in ice-cold 1x PBS and then collected by scraping in 1x PBS. The cells were incubated in Lysis Buffer 1 [50 mM HEPEs pH 7.5, 140 mM NaCl, 1mM EDTA, 10% (v/v) glycerol, 0.5% (v/v) IGEPAL CA-630, 0.25% (v/v) Triton X-100, 1x protease inhibitor cocktail] for 10 minutes on ice. The resulting nuclei were pelleted and washed with Lysis Buffer 2 (10 mM Tris pH 8.0, 200 mM NaCl, 1mM EDTA, 0.5 mM EGTA, 1x protease inhibitor cocktail) and pelleted. The nuclei were then lysed using Lysis Buffer 3 [10 mM Tris pH 8.0, 100 mM NaCl, 1 mM EDTA, 0.5 mM EGTA, 0.1% (w/v) sodium deoxycholate, 0.5% (v/v) N-lauroylsarcosine, 1x protease inhibitor cocktail] and sonicated for 10 minutes (30 sec on/30 sec off) using the Diagenode Pico sonicator. The sheared chromatin was cleared of debris by applying Triton X-100 to a final concentration of 1% (v/v), followed by centrifugation at >20,000 x g for 10 minutes at 4°C. The clarified chromatin samples were then applied to the antibody-loaded beads overnight at 4°C with gentle mixing. The chromatin-immune complexes on the beads were washed seven times with RIPA buffer, followed by two washes in 100 mM ammonium hydrogen carbonate (AMBIC) solution to remove the detergents. The samples were de-crosslinked by boiling at 4°C for 10 minutes.

#### Mass spectrometry

After reduction and alkylation with tris(2-carboxyethyl) phosphine hydrochloride (TCEP; Sigma, 646547-10X1ML) and iodoacetamide (IAA; Sigma, I1149-5G), the samples were digested overnight with trypsin (Pierce, 90057). After solid-phase extraction cleanup with an Oasis HLB µelution plate (Waters), the resulting peptides were reconstituted in 2% (v/v) acetonitrile (ACN) and 0.1% trifluoroacetic acid in water to a concentration of about 0.5 μg/μL. Two μL of each sample were injected onto a Q Exactive HF mass spectrometer coupled to an Ultimate 3000 RSLC-Nano liquid chromatography system with a 75 um i.d., 15-cm long EasySpray column (Thermo Scientific, ES75150PN) and eluted with a gradient from 0-28% Buffer B over 90 min. Buffer A contained 2% (v/v) ACN and 0.1% formic acid in water, and Buffer B contained 80% (v/v) ACN, 10% (v/v) trifluoroethanol, and 0.1% formic acid in water. The mass spectrometer operated in positive ion mode with a source voltage of 2.5 kV and an ion transfer tube temperature of 300°C. MS scans were acquired at 120,000 resolution in the Orbitrap and up to 20 MS/MS spectra were obtained in the ion trap for each full spectrum acquired using higher-energy collisional dissociation (HCD) for ions with charges 2-8. Dynamic exclusion was set for 20 s after an ion was selected for fragmentation.

#### Mass spectrometry data analyses

Raw MS data files were analyzed using Proteome Discoverer v3.0 (Thermo Fisher, RRID:SCR_014477), with peptide identification performed using Sequest HT searching against the human reviewed protein database from UniProt. Fragment and precursor tolerances of 10 ppm and 0.02 Da were specified, and three missed cleavages were allowed. Carbamidomethylation of Cys was set as a fixed modification and oxidation of Met was set as a variable modification. The false-discovery rate (FDR) cutoff was 1% for all peptides. Peptide peak intensities were summed for all peptides matched to a protein for protein quantitation.

Significance of enrichment differences between the E2 treatment and vehicle control values were calculated for each identified protein using a Student’s t-test. Proteins with p-values < 0.05 were used for further analysis. Relative abundances were calculated by dividing the abundance values found from the ERα immunoprecipitations with the corresponding IgG control precipitations. The E2 dose at the which relative abundance was maximal was calculated using the max() function in R, and each identified protein was categorized as such. The DAVID online tool (Dennis et al., 2003) was used to determine enriched gene ontology terms for these categories (see below).

### Gene ontology and gene set enrichment analyses

#### Gene ontology analysis

Gene Ontology analysis of the RNA-seq data was conducted using the Database for Annotation, Visualization, and Integrated Discovery (DAVID) online tool (Dennis et al., 2003) and the ontological terms were ranked based on enrichment scores.

#### Kyoto encyclopedia of genes and genomes (KEGG) analysis

Gene set enrichment analysis was performed on RNA-seq differential expression data using KEGG analysis (Kanehisa et al., 2023) with the clusterProfiler R package using a p-value cutoff of < 0.05.

### DNA sequence motif analyses

#### HOMER analysis

De novo motif analysis was performed using the HOMER findmotifGenome.pl command line (Heinz et al., 2010), and known motifs were used for subsequent analyses. For motif analysis around ChIP-seq peaks, these analyses were performed on a 200 bp region surrounding the peak summits. For analyses performed on open chromatin regions, ATAC-seq data was used and the analyses were done within the peak range (-d given). Gene logos were obtained from the corresponding motifs using the JASPAR database (Sandelin et al., 2004).

#### Find individual motif occurrences (FIMO) analysis

FIMO (Grant et al., 2011) was used to identify motifs within initially opened and newly opened regions of individual gene regions. Regions of interest were identified based on peak occurrences in ATAC-seq data. DNA sequences within these regions were uploaded as a .fasta file to the FIMO web tool in the MEME Suite 5.5.8 (Bailey et al., 2009) using a p-value cutoff of < 0.005.

### Survival plots

Kaplan-Meier curves for overall survival analyses were made using the GEPIA2 open-source database containing RNA-seq gene expression data (Tang et al., 2019). The data were first sorted for all breast cancer patients, followed by the luminal A molecular subtype, and finally by gene signature. A quartile cutoff was applied, and patients falling under a 95% confidence interval was plotted over a 5-year period.

### Statistical analyses

All genomic analyses were performed at a minimum of two individual biological replicates. Statistical analysis for the genomic experiments was performed using standard genomic statistical tests. All qPCR-based experiments had a minimum of three independent biological replicates. All statistical tests used and their corresponding p-values are provided in the figure legends.

## Acknowledgements

The authors would like to thank the following: (1) Dr. Dan Huang for helpful feedback and assistance with editing the manuscript and figures, (2) Sneh Koul for additional computational support, (3) Dr. Yoon Jung Kim from UT Southwestern’s Children’s Research Institute Sequencing Core, (4) Dr. Andrew Lemoff from UT Southwestern’s Proteomics Core, (5) Vanessa Schmid from UT Southwestern’s McDermott Center Next Generation Sequencing Core, and (6) members of the Kraus lab for continued input and feedback on this project.

## Funding

This work was supported by a grant from the NIH/NIDDK (R01 DK058110) and funds from the Cecil H. and Ida Green Center for Reproductive Biology Sciences Endowment to W.L.K.

## Author Contributions

H.B.K. and W.L.K. conceived and developed this project, designed the experiments, and oversaw their execution. C.V.C provided intellectual input and H.B.K. preformed all of the wet lab experiments. H.B.K. and T.N. performed the computational analyses of the genomic and proteomic data. H.B.K. prepared the initial drafts of figures and text, with assistance from T.N. and W.L.K. The figures and text were edited and finalized by W.L.K and C.V.C. W.L.K. secured funding to support this project, and provided intellectual support and overall leadership for the work, with assistance from C.V.C.

## Competing Interests

The authors have no competing interests to declare.

## Data and Materials

Additional details for the protocols and data analyses, raw data from the qPCR-based assays, and experimental materials are available upon request from the corresponding author. The raw genomic can be accessed from NCBI’s Gene Expression Omnibus (GEO) database using the following accession numbers: GSE298771 (RNA-seq), GSE298767 (ChIP-seq), GSE298769 (ATAC-seq), and GSE298770 (PRO-seq) under the super series accession number GSE298773. The raw RIME proteomics data can be accessed from the Mass Spectrometry Interactive Virtual Environment (MassIVE) data base using accession number MSV000098072.

## Supplementary Figure Legends

**Supplementary Figure S1. Overlap of regulated genes sets and gene set enrichment analyses (GSEA).**

**(A and B)** Venn diagram showing overlap of differentially (A) upregulated or (B) downregulated genes in the pM responsive (*green*; 1 pM, 10 pM, and 100 pM E2) and nM responsive groups (*orange*; 1 nM, 10 nM and 100 nM E2). p < 0.05; log_2_(FC) > 0.6 for upregulated genes and log_2_(FC) < −0.6 for downregulated genes.

**(C and D)** GSEA reveals pathways that are differentially enriched between pM responsive and nM responsive gene sets. KEGG analysis of genes expressed within (C) pM responsive and (D) nM responsive gene sets from RNA-seq data. Significance values shown with shading and number of terms within each group shown with size of data points.

**Supplementary Figure S2. Genomic analyses across a range of E2 concentrations.**

**(A)** Heatmap showing ERα chromatin occupancy as determined by ChIP-seq for a range of E2 concentrations (vehicle, 10 pM, 100 pM, 1 nM, and 10 nM). The data are centered on the peak summits and are shown for a ± 5 kb region around the summits.

**(B)** Euler plot showing overlaps of ERα ChIP-seq peaks for a range of E2 concentrations.

**(C and D)** Chromatin accessibility by ATAC-seq (C) and H3K27ac enrichment by ChIP-seq (D) determined for a range of E2 concentrations. The data are centered on the peak summits and are shown for a ± 5 kb region around the summits.

**Supplementary Figure S3. Dose-dependent binding of ERα, FOXA1, or GATA3 to chromatin.**

**(A)** Chromatin enrichment assays showing the levels of ERα, FOXA1, and GATA3 in a chromatin fraction from MCF-7 cells treated with vehicle (DMSO), 10 pM, or 10 nM E2 for 30 minutes after siRNA knockdown of *FOXA1*, *GATA3*, or a non-targeting control. The proteins were detected by Western blotting. Histone H3 was used as a loading control.

**(B)** Table showing the total number of FOXA1 or GATA3 peaks determined by ChIP-seq after vehicle (DMSO), pM (10 pM E2), and nM (10 nM E2) treatment. The number of peaks that overlap with ERα binding and were used for subsequent analyses are shown in the final column.

**(C)** Venn diagrams showing overlapping peaks between vehicle (DMSO) and nM (10 nM E2) treatment conditions for FOXA1 (*green*) and GATA3 (*orange*) at ERα binding sites.

**(D)** Boxplot showing the ratio of the chromatin occupancies for GATA3 and FOXA1 ± 250 bp around ERα peak summits for pM responsive and nM responsive ERBS. Wilcoxon Rank Sum test, **** = p < 5 × 10^-15^.

**Supplementary Figure S4. Dose-dependent enrichment of c-Fos (AP-1) motifs near ERα binding sites.**

**(A)** JASPAR gene logo of AP-1 motifs found ± 200 bp around ERBSs (*top*) and bar graph showing the abundance of motif occurrences around ERBSs established over a range of E2 concentrations, normalized to their respective maximum values (*bottom*).

**(B)** Browser tracks showing ERα ChIP-seq data from MCF-7 cells at two different concentrations of E2 aligned with c-Fos ChIP-seq data for representative pM responsive (*XBP1*) and nM responsive (*PGR*) genes. The dashed box highlights a region (*bottom*) that shows overlap of ERα and c-Fos binding in the nM responsive gene.

**(C)** Euler diagrams showing overlaps of ERα ChIP-seq peaks across a range of different E2 treatment concentrations (10 pM to 10 nM) with c-Fos ChIP-seq peaks.

**Supplementary Figure S5. Summary of sequencing analysis at pM responsive and nM responsive ERα enhancers and genes.**

**(A and B)** Line graphs showing relative chromatin accessibility (ATAC-seq), transcription (PRO-seq), and ERα chromatin occupancy (ChIP-seq) around ERα enhancers (*left*) and gene expression (RNA-seq), transcription (PRO-seq), and RNA Pol II pausing (pausing index from PRO-seq) at the corresponding target genes (*right*). Analysis was done for pM responsive (A) and nM responsive (B) ERα enhancers and genes. All values were normalized to their respective maximums.

**Supplementary Figure S6. ERα-RIME proteomic analysis showing protein abundance patterns for pM and nM E2 treatments**

**(A)** Coverage map from RIME mass spectrometry data for ERα using the Sequence Coverage Visualizer web application.

**(B and C)** Line graphs showing ERα-RIME data expressed as relative abundances for proteins with maximum enrichment at 10 pM E2 (B, n = 275) or 10 nM E2 (C, n = 276). Gray lines show the enrichment values from individual proteins, while the colored lines show the average.

(**D and E**) GO analysis showing the number of proteins and significance values for the top 5 terms from ERα-RIME data for proteins with maximum enrichment at 10 pM (D) or 10 nM (E) E2 treatment.

**(F)** Boxplots showing the relative abundance of coregulators that associate with chromatin-bound ERα as determined by RIME. The abundances were normalized to the respective IgG controls. Representative proteins with maximum abundance at 10 pM E2 (p300 and NCOA3/SRC3) or 10 nM E2 (NCOA2/SRC2) are shown. Bars marked with different letters are significantly different from each other. Student’s t-test, p < 0.005.

**Supplementary Figure S7. Overlap of proteins identified by ERα-RIME and FERM eRNA pulldown.**

Venn diagram showing the overlap between proteins identified by ERα-RIME compared to those identified by eRNA FERM element pulldown (*left*). Table showing overlapping proteins categorized by the E2 dose where maximum abundance was observed as determined by ERα-RIME (*right*).

## Notes

### Competing Interest Statement

The authors have declared no competing interest.

