## Supplementary Figures S1-S7 for "Estrogen Receptor Enhancers Sensitive to Low Doses of Hormone Specify Distinct Molecular and Biological Outcomes"

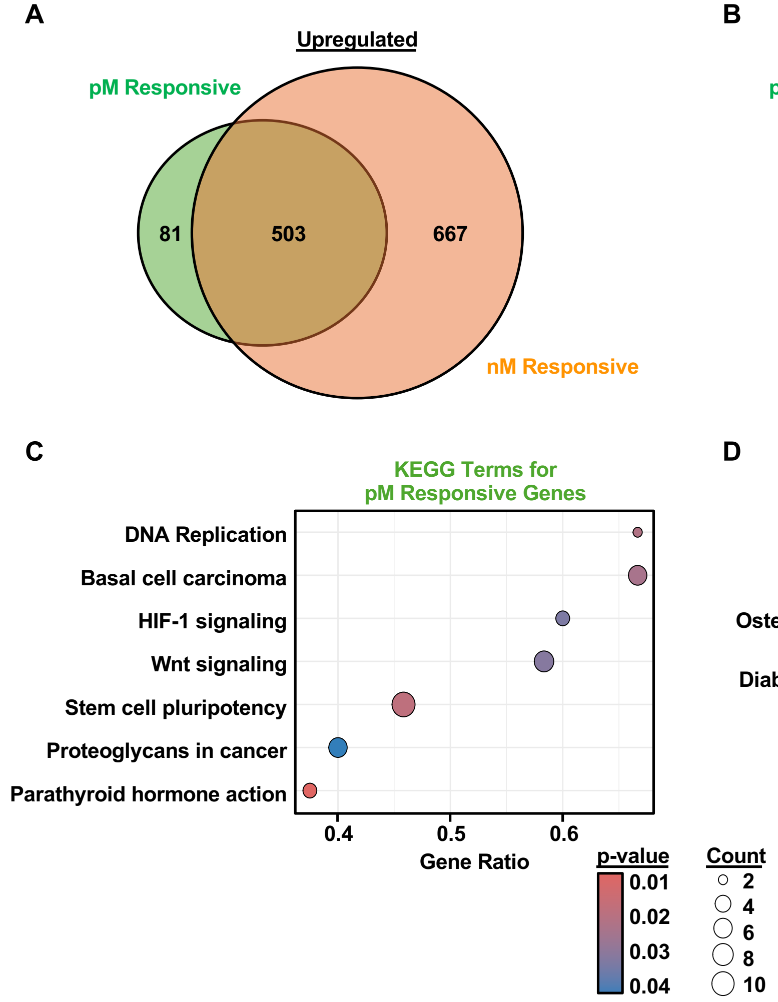
A
Upregulated
pM Responsive
81
503
667
nM Responsive
B
Downregulated
pM Responsive
78
287
664
nM Responsive
C
KEGG Terms for
pM Responsive Genes
DNA Replication
Basal cell carcinoma
HIF-1 signaling
Wnt signaling
Stem cell pluripotency
Proteoglycans in cancer
Parathyroid hormone action
0.4
0.5
0.6
Gene Ratio
D
KEGG Terms for
nM Responsive Genes
HSV1 infection
Cellular senescence
FoxO signaling
Osteoclast differentiation
T-Cell leukemia
Diabetic cardiomyopathy
Insulin signaling
MAPK signaling
Tight junction
HPV infection
0.4
0.6
0.8
Gene Ratio
p-value
0.01
0.02
0.03
0.04
Count
2
4
6
8
10

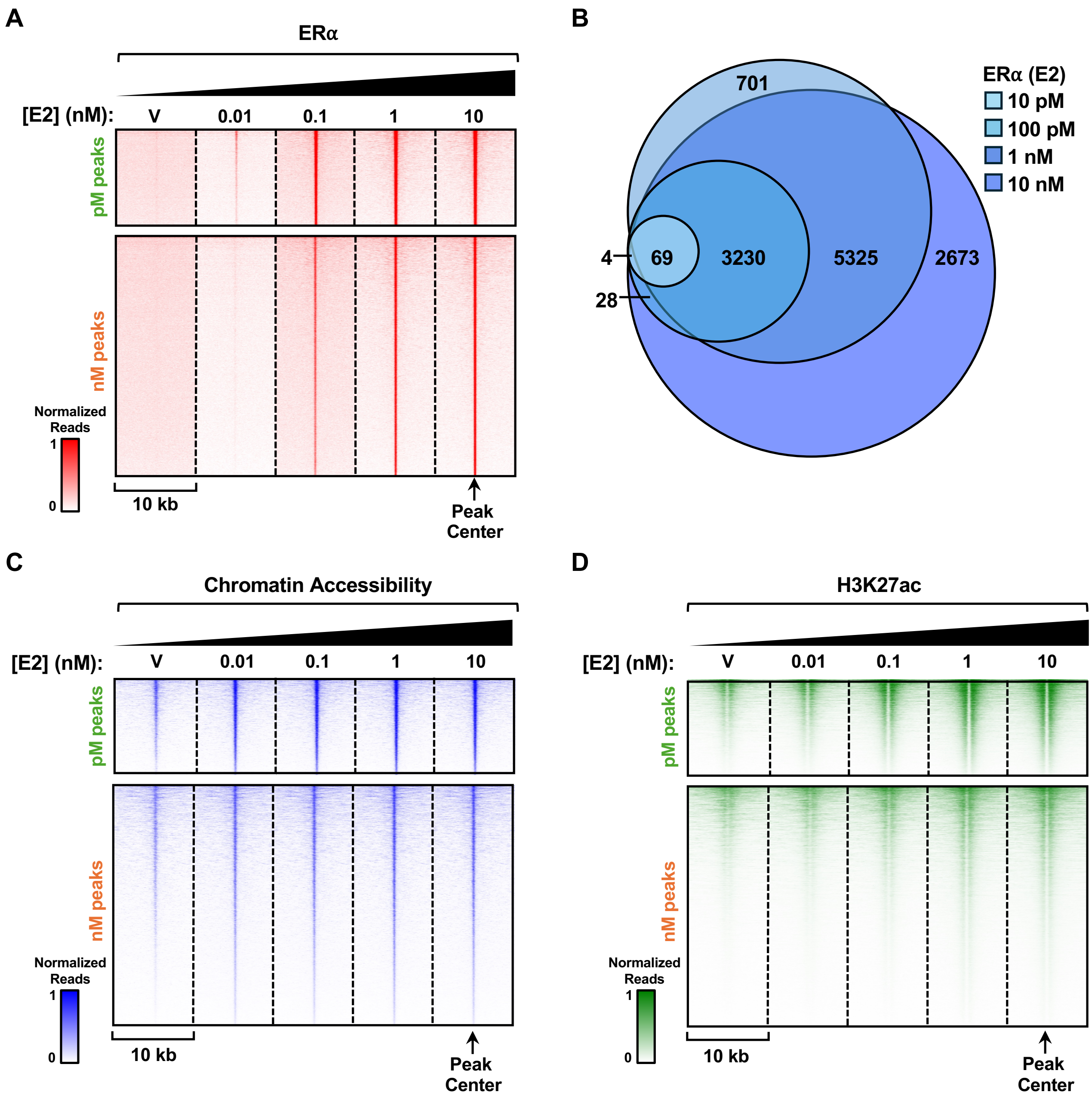


Kim *et al.* (2025) – Supplementary Figure S2

A

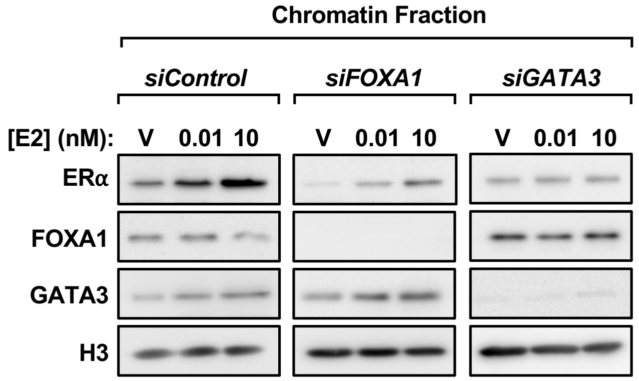


B

| | Total Number of Peaks | | | |
|---|---|---|---|---|
| ChIP-seq | Vehicle | 10 pM E2 | 10 nM E2 | Overlap with ERα |
| FOXA1 | 49,027 | 51,690 | 39,502 | 2,658 |
| GATA | 2,014 | 8,368 | 7,325 | 4,138 |

C

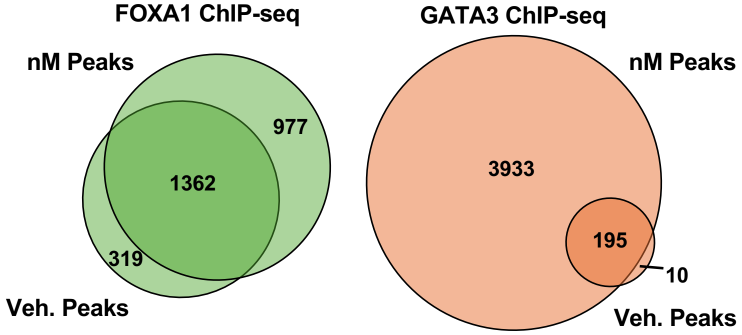


D

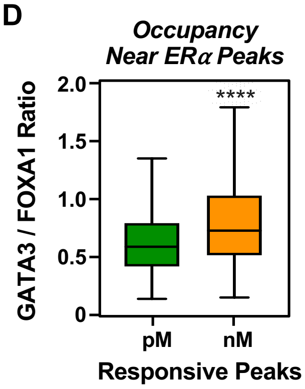

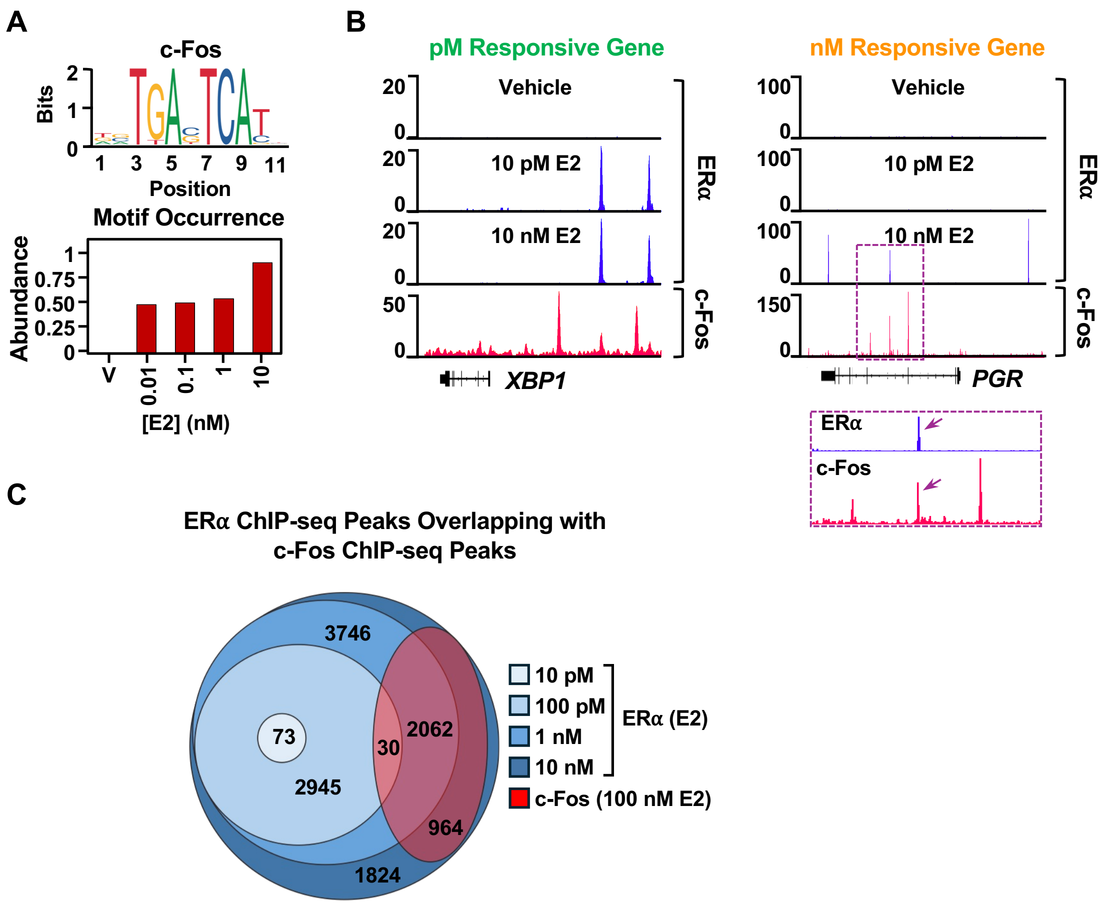


Kim *et al.* (2025) – Supplementary Figure S4

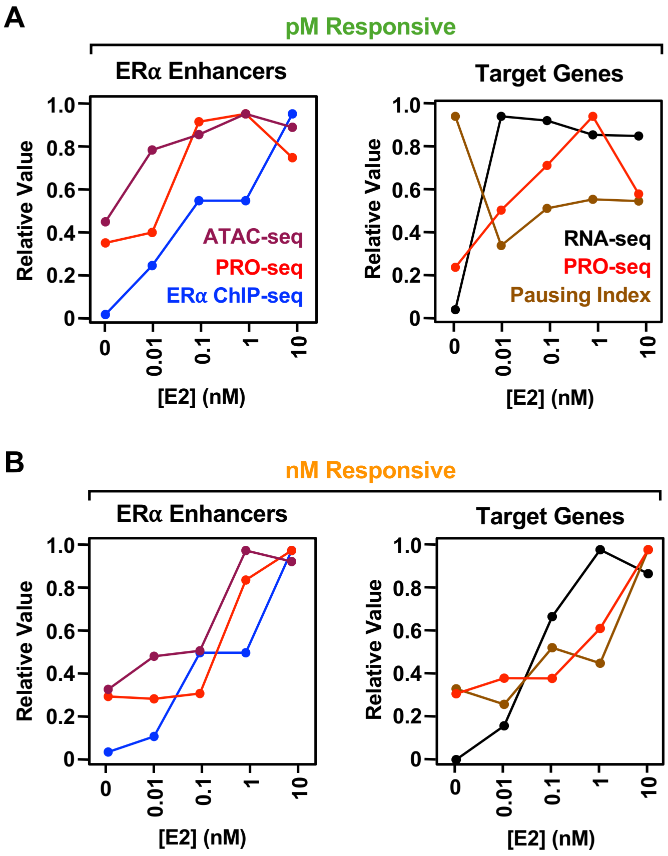
A
pM Responsive
ERα Enhancers
Target Genes
Relative Value
1.0
0.8
0.6
0.4
0.2
0
ATAC-seq
PRO-seq
ERα ChIP-seq
RNA-seq
PRO-seq
Pausing Index
0
0.01
0.1
1
10
[E2] (nM)
B
nM Responsive
ERα Enhancers
Target Genes
Relative Value
[E2] (nM)

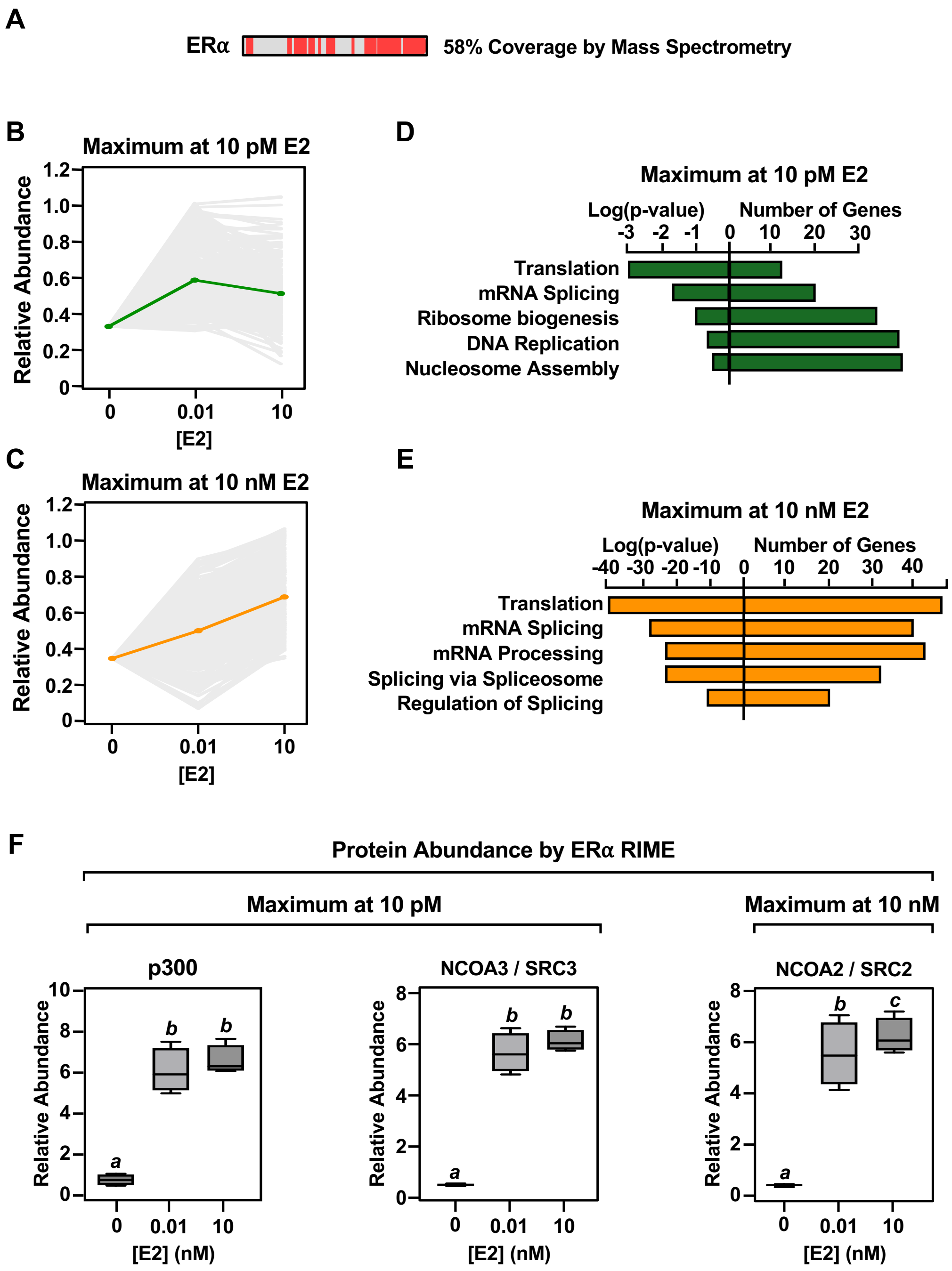


Kim *et al.* (2025) – Supplementary Figure S6

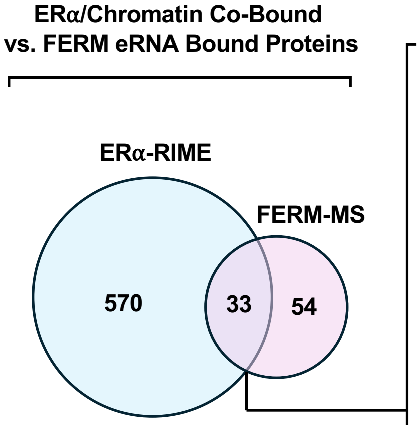


**Overlapped by Dosage**

| pM | nM |
|---|---|
| DHX15 | HNRNPH2 |
| SNRPB | SNRPA1 |
| RUVBL2 | HNRNPH1 |
| NPM1 | DPF2 |
| AP1M2 | SYNCRIP |
| DYNLL1 | YY1 |
| RACK1 | HNRNPC |
| TUBB | TUFM |
| RBBP7 | PCBP2 |
| COPE | SRSF6 |
| UPF1 | CAPZB |
| THRAP3 | PCBP1 |
| BCLAF1 | HNRNPF |
| PHF6 | HNRNPD |
| SLC25A3 | ILF2 |
| PHF5A | AGR2 |
| SLC25A6 | |
